# LiftPose3D, a deep learning-based approach for transforming 2D to 3D pose in laboratory animals

**DOI:** 10.1101/2020.09.18.292680

**Authors:** Adam Gosztolai, Semih Günel, Victor Lobato Ríos, Marco Pietro Abrate, Daniel Morales, Helge Rhodin, Pascal Fua, Pavan Ramdya

## Abstract

Markerless 3D pose estimation has become an indispensable tool for kinematic studies of laboratory animals. Most current methods recover 3D pose by multi-view triangulation of deep network-based 2D pose estimates. However, triangulation requires multiple, synchronized cameras and elaborate calibration protocols that hinder its widespread adoption in laboratory studies. Here, we describe LiftPose3D, a deep network-based method that overcomes these barriers by reconstructing 3D poses from a single 2D camera view. We illustrate LiftPose3D’s versatility by applying it to multiple experimental systems using flies, mice, rats, and macaque monkeys and in circumstances where 3D triangulation is impractical or impossible. Our framework achieves accurate lifting for stereotyped and non-stereotyped behaviors from different camera angles. Thus, LiftPose3D permits high-quality 3D pose estimation in the absence of complex camera arrays, tedious calibration procedures, and despite occluded body parts in freely behaving animals.

## 1 Introduction

To identify how actions arise from neural circuit dynamics, one must first make accurate measurements of behavior in laboratory experiments. Paired with new methods for recording neuronal populations in behaving animals [1–4], recent innovations in 3-dimensional (3D) pose estimation promise to accelerate the discovery of fundamental neural control principles. 3D pose estimation is typically accomplished by triangulating 2-dimensional (2D) poses acquired using multiple camera views and deep network-based markerless pose tracking algorithms [5–13]. Notably, triangulation requires that every tracked keypoint, be it a joint or other body feature, be visible from at least two synchronized cameras [14] and that each camera be calibrated. This can be done by hand [15, 16] or, by solving a non-convex optimization problem [7]. These expectations are high and often difficult to meet, particularly in space-constrained experimental systems that also house sensory stimulation devices [1, 2, 17]. When untethered and freely behaving animals, such as fur-covered rodents [18], are observed under these conditions, some limb keypoints are often intermittently occluded in some camera views, meaning that 3D triangulation may be impossible for these keypoints.

Because of this, most animal studies have favored simple and higher throughput 2D pose estimation approaches using only one camera [5, 6, 10, 19–21]. Nevertheless, 3D poses are still desirable, among other reasons because they eliminate the otherwise present camera-angle dependence of behavioral analyses based on 2D poses [7]. Computer vision research on human pose estimation has long been interested in “lifting” 2D poses, that is, recovering 3D poses by regression to a ground truth dataset of 3D poses [22–25] but only recently have deep learning-based methods achieved high accuracy [26–38]. However, these techniques have not yet been adapted to laboratory animal studies due to the above mentioned challenges of acquiring large and diverse training datasets of behaving animals. Additionally, in some experiments, 3D ground truth data is completely missing. This prohibits training a lifting network and creates the need to generalize pre-trained lifting networks across experimental systems.

Here, we introduce LiftPose3D, a deep learning-based tool for frame-by-frame 3D pose estimation of tethered and freely behaving laboratory animals from a single camera view. Our method relies on a neural network architecture initially designed to lift human poses [34]. Due to its simplicity, this network does not require temporal information or a skeletal graph. Hence, it generalizes easily. We develop data transformations and network training augmentation methods that enable accurate 3D pose estimation across a wide range of animals, camera angles, experimental systems, and complex behaviors using relatively little data. Our findings are as follows:

1. We show that a library of 3D poses can be used to train a network to lift 3D poses from a single cameraïs annotated 2D poses. We impose minimal constraints on the camera hardware and do not require *a priori* knowledge about camera position. Consequently, our method does not require prior camera calibration.
2. We demonstrate that alignment of animal poses into the same reference frame allows the network to learn relationships between pose keypoints. We use this to (i) predict complete 3D poses in freely behaving animals despite occlusions and to (ii) correct outliers in ground truth data.
3. By varying the bone lengths of pose skeletons during training, our method gains robustness to large variations in animal body proportions.
4. We find that pose differences between experimental domains are mostly linear and that pre-trained LiftPose3D networks can be adapted to generalize using a linear domain adaptation technique.

We illustrate these findings in several experimental scenarios. First, for tethered adult *Drosophila* [7] and freely behaving macaque monkeys [8], we use LiftPose3D to reduce the number of cameras required for 3D pose estimation, often to a single camera, and relax constraints on camera placement. We make these pretrained networks and our code publicly available to be used for new experiments in other laboratories. Second, for freely behaving *Drosophila*, mice [18], and rats [39], LiftPose3D can obtain 3D poses despite occlusions. Finally, using linear domain adaptation, pretrained LiftPose3D networks can be used to predict realistic 3D poses from different experimental systems viewing *Drosophila* behaviors ventrally with a single camera. This technique allows us to effectively resurrect old data for new kinds of kinematic analyses [20]. To reduce the entry barrier for users interested in obtaining 3D pose data in this manner, we explain how to construct a cheap and reliable hardware system that we call a *Drosophila* “LiftPose3D station.”

## 2 Results

### 2.1 Theoretical basis for LiftPose3D

If a keypoint *j* of interest is visible from at least two cameras, with corresponding 2D coordinates **x**_*c,j*_ in camera *c* and camera parameters (extrinsic and intrinsic matrices, see Materials and Methods for details), then its 3D coordinates **X**_*j*_ in a global world reference frame can be obtained by triangulation. Here we use triangulated 3D positions as ground truth with which to assess the accuracy of LiftPose3D, a method that focuses on lifting 3D poses from a single camera. Rather than considering keypoints independently, our goal is to predict the coordinates of *n* keypoints **X** = (**X**_1_, …, **X**_*n*_)—the 3D pose— from their respective 2D coordinates **x**_*c*_ = (**x**_*c*,1_, …, **x**_*c,n*_) viewed from a camera *c*. By considering all keypoints simultaneously, our method hinges upon learning spatial relationships between them in the context of animal poses. Moreover, we seek to impose minimal constraints on camera *c* meaning that its parameters need not be known (e.g., see **Figure 1**A, illustrating six fixed cameras).

**Figure 1:**
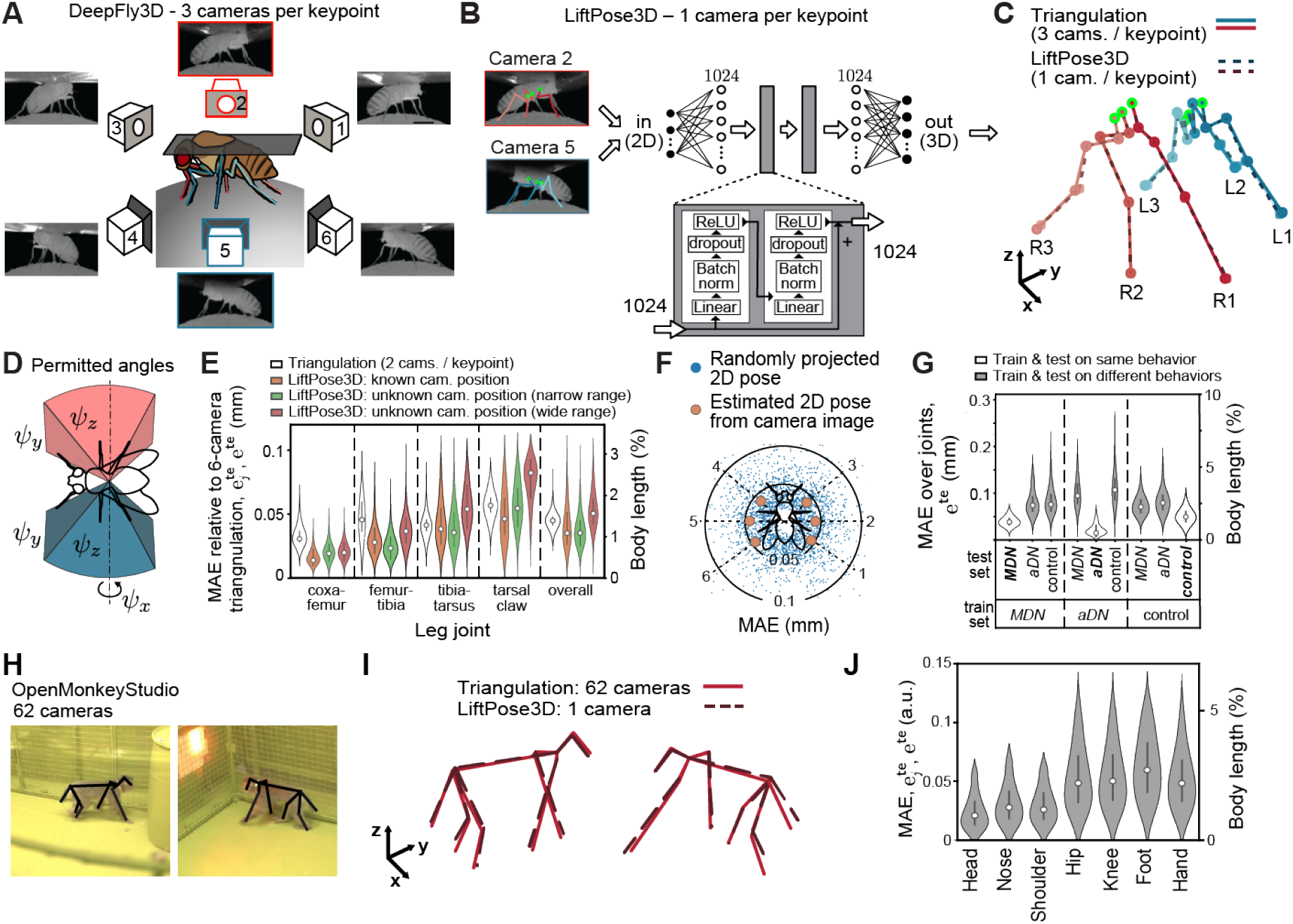
LiftPose3D predicts 3D pose with fewer cameras and flexible camera positioning. **A** Ground truth 3D poses of tethered *Drosophila* are triangulated using six camera views (3 cameras per keypoint). LiftPose3D predicts 3D poses using only two cameras (red and blue, 1 camera per keypoint). **B** As inputs, LiftPose3D takes deep network-derived 2D poses for 15 joints per camera (red and blue). The coordinates of the 2D poses are considered relative to a set of root joints (green). The inputs are scaled to 1024 dimensions by an affine layer, passed twice through the main processing unit (gray rectangle). The main processing unit consists of two fully-connected layers of 1024 dimensions wrapped by a skip connection, consisting of batch norm, dropout and ReLU. **C** The output of the network are 3D poses for the left (blue) and right (red) body halves, which are compared with the ground truth 3D poses obtained from triangulation. Limbs are labeled according to left/right and front (1), mid(2), or hind (3) position. **D** Permitted camera placements. By making virtual camera projections of the 3D pose within angles *ψ*_*z*_, *ψ*_*y*_, *ψ*_*x*_ (representing ordered yaw, roll, pitch rotations) LiftPose3D can be be trained to lift from cameras placed at any angle. **E** Error of lifted 3D poses relative to triangulation using three cameras per keypoint. Violin plots show the triangulation error using the theoretical minimum of 2 cameras per keypoint (white), test error for a network trained with known camera parameters (orange) and two angle-invariant networks with narrow (green) and wide ranges (red). **F** Error of lifted 3D poses at different virtual camera orientations of the wide-range angle-invariant lifter network and a network with known camera parameters. Blue dots represent lifting errors for a given projected 2D pose. Orange circles represent averages over the test dataset from a given camera. **G** Error of estimated 3D poses for a LiftPose3D network trained and tested on different combinations of data containing flies performing optogenetically-induced backward walking (*MDN*, left), antennal grooming (*aDN*, middle), or spontaneous (unstimulated) behaviors (*PR*, right). **H** Two representative images from the OpenMonkeyStudio dataset. 2D poses are superimposed (black). **I** 3D poses obtained by triangulating up to 62 cameras (red lines) or using a single camera and LiftPose3D (dashed black lines). **J** Distribution of absolute errors for different body parts with respect to total body length. Violin plots represent Gaussian kernel density estimates with bandwidth 0.5, truncated at the 99th percentile and superimposed with the median (gray dot), 25th, and 50th percentiles (black line).

The basis of LiftPose3D is to estimate the 3D pose by learning a nonlinear mapping between triangulated ground truth 3D poses and corresponding 2D poses. Formally, this operation is encoded in a *lifting* function *f* mapping a 2D pose from any camera *c* to their corresponding 3D pose in camera-centered coordinates, **Y**_*c*_ = *f* (**x**_*c*_), and a camera transformation *ϕ*_*c*_, encoding a rotation and translation operation (see Eq. (2) in the Materials and Methods), mapping from camera-centered coordinates to world coordinates 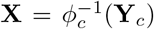. The lifting function *f* can be approximated by a deep neural network *F* (**x**_*c*_; Θ), where Θ represents the network weights controlling the behavior of *F*. In a specific application, Θ are trained by minimizing the discrepancy between 3D poses predicted by lifting from any camera and ground truth 3D poses,

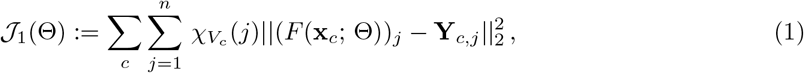

where 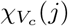 is an indicator function of the set *V*_*c*_ of visible points from camera *c*. For *F* (**x**_*c*_; Θ), we adapt a network architecture from [34] composed of fully connected layers regularized by batch-norm and dropout [40] and linked with skip connections (**Figure 1B**). This network has been previously developed for human-pose estimation to be trained on approximately 10^6^ fully annotated 2D-3D human pose pairs for many different behaviors. By constrast, we will demonstrate that training augmentation methods allow this network to (i) work with a vastly smaller training dataset (between 10^3^-10^4^ poses acquired automatically using 2D pose estimation approaches [6, 7]), (ii) predict 3D poses from a single camera view at arbitrary angles, (iii) be trained with only partially annotated ground truth 3D poses suffering from occlusions, and (iv) generalize a single pretrained network across experimental systems and domains through linear domain adaptation.

Note that our setup in Eq. (1) implicitly assumes that the network learns two operations: lifting the 2D pose **x**_*c*_ to camera-centered 3D coordinates **Y**_*c*_ by predicting the depth component of the pose, and learning perspective effects encoded in the animal-to-camera distance and the intrinsic camera matrix (see Eqs. (2)–(5) in Materials and Methods). Notably, the intrinsic camera matrix is camera-specific, suggesting that a trained network can only lift poses from cameras used during training and that application to new settings with strong perspective effects (short focal lengths) may require camera calibration. We will show that this is not necessarily the case and that one can generalize pre-trained networks to new settings by weakening the perspective effects. This can be accomplished by either using a large focal length camera, or by increasing the animal-to-camera distance and normalizing the scale of 2D poses [41] (see Materials and Methods). We will demonstrate that a weak perspective assumption can, in many practical scenarios, enable lifting 2D poses from different cameras without calibration. As well illustrate next, these contributions enable 3D pose estimation in otherwise inaccessible experimental scenarios.

### 2.2 Predicting 3D pose with fewer cameras, flexible positioning, and diverse camera hardware

To illustrate how LiftPose3D can simplify 3D pose acquisition, we considered a previously published tethered adult *Drosophila* dataset [7]. This dataset is representative of current laboratory practice of obtaining 3D poses by triangulation of multiple, synchronized camera views per keypoint [7,16]. Here, 15 keypoints on each lateral side of the animal (**Figure 1A**) were annotated by DeepFly3D [7] and triangulated from three camera views. Using LiftPose3D, we aimed to reduce the number of cameras needed for 3D pose estimation to two, i.e., one camera per keypoint, where triangulation is not possible (**Figure 1B**). Furthermore, the requirement to know the cameras’ positions for calibration purposes can be eliminated for long focal length cameras.

We envisioned that, using this tethered *Drosophila* dataset [7] as a 3D pose library, we might train a LiftPose3D network to be directly applied to other experiments. To achieve this goal, we needed to ensure that the output of LiftPose3D would be independent of any translations of input 2D poses, perspective effects, and the placement of the camera. First, to achieve translation invariance, we predicted the keypoints of the respective legs relative to a set of six “root” keypoints, which we chose to be the immobile thorax-coxa joints (green circles, **Figure 1B**). Second, to factor out perspective effects, we assumed that the focal length of the camera and the animal-to-camera distance are either known or that one of them is large enough to assume weak perspective effects. In the latter case, we normalized 2D input poses by their Frobenius norm at both training and test times. Third, to obtain camera-angle invariance, we parametrized the possible camera orientations by Euler angles *ψ*_*z*_, *ψ*_*y*_, *ψ*_*x*_ representing ordered rotations around the *z, y* and *x* axes of a coordinate system centered around the fly (**Figure 1D**). During training, we took as outputs ∼2.5 × 10^4^ 3D poses obtained from three-camera triangulation and obtained input 2D poses by randomly projecting to virtual camera planes within specified Euler angle ranges. We trained a “narrow angle-range” network with Euler angles around a known camera location (*ψ*_*z*_ = ±10°, *ψ*_*y*_ = ±5°, *ψ*_*x*_ = ±5°), or a “wide angle-range” network covering all camera locations around the meridian (*ψ*_*z*_ = ± 180°, *ψ*_*y*_ = ±5°, *ψ*_*x*_ = ±5°). Importantly, beyond weak perspective, no assumption was made about the camera positioning and lens focal lengths during training. As a baseline scenario where the camera parameters are known, we also trained a network using 3D poses as outputs and 2D poses obtained from DeepFly3D-annotated images as inputs. We tested each LiftPose3D network by predicting ∼3.6 × 10^3^ triangulated 3D poses from two independent animals and software-annotated 2D poses from side camera images (**Figure 1B**; cameras 2 and 5). We evaluted the networks’ predictions relative to the triangulated ground truth by computing the mean absolute error (MAE), 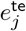, for each joint *j* as well as the MAE across all joints 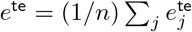

We found that LiftPose3D could predict 3D poses using only one camera per side (**Figure 1C**). When we trained and tested the network using poses from the same set of cameras, the accuracy was at least as good as from triangulation using two cameras per keypoint (**Figure 1E**, white). More surprisingly, the accuracy did not suffer for the narrow angle-range network (**Figure 1E**, green), which was trained using virtual 2D projections (rather than true 2D pose estimates), and for which the intrinsic camera parameters were unknown. For the wide angle-range network spanning the full 360°(**Figure 1E**, red), the accuracy remained excellent. This is illustrated in videos of lifted 2D poses from animals that were optogenetically induced to walk backwards (**Video 1**) or groom their antennae (**Video 2**). It was also true for animals generating spontaneous, irregular limb movements, demonstrating that that lifting can be performed as well for complex, non-stereotyped movements (**Video 3**). Although accuracy was high for all keypoints, the MAE progressively increased from the proximal to distal joints. This is expected because the network predicts joint coordinates with respect to the thorax-coxa root joints and nearby, proximal joints move within a smaller kinematic volume. By contrast, triangulation obtains the 3D coordinates for each keypoint independently and, consequently, its error depends only on the accuracy of underlying 2D annotations. Next, to assess the camera-angle dependence of the test error for the wide angle-range network, we either generated virtual projections on the meridian of the unit sphere, or lifted 2D poses from each of the six known cameras (**Figure 1F**). The MAE was low (*<* 0.05 mm) for all camera arrangements with no clear camera-angle dependence. Since our angle-invariant lifter networks are trained using virtual projections, they make no assumptions about camera hardware or positioning. These results imply that our pretrained networks can provide a simple yet accurate means of obtaining 3D poses for tethered *Drosophila* systems in other laboratories.

We predicted that lifting accuracy would also depend on the degree of overlap between behaviors found in the training and test datasets. This is an important dimension to explore, given the relatively small amounts of data available from laboratory experiments. The tethered *Drosophila* dataset contained optogenetically-induced behaviors like antennal grooming (*aDN*), and backward walking (*MDN*), as well as spontaneously-generated behaviors like forward walking. We trained LiftPose3D using poses from only one of these behaviors (eliminating frames where the animal was resting), while keeping the amount of training data (2.5 ×10^4^ poses) fixed, and evaluated the network performance on all three behaviors. As expected, the MAE was higher when test data included untrained optogenetically-induced and spontaneously-generated control behaviors (*PR*) than for test data with the same behaviors as in the training data (**Figure 1G**). Furthermore, a network trained on all three behaviors showed comparable or lower MAE (**Figure 1E**, orange) than networks trained and tested on the same specific behavior (**Figure 1G**). Thus, a behaviorally diverse training dataset can be expected to lift 3D poses with more accuracy than a dataset with fewer behaviors.

Having accurate 3D poses confers several advantages, including eliminating artifactual camera angle-dependencies in downstream analyses such as behavioral clustering [7]. To further illustrate the added benefit of 3D poses over 2D poses, we illustrate joint angles during forward walking from lifted 3D poses (*α, β, γ, ω*, **Figure S1**, red), from 3D triangulated ground truth poses (**Figure S1**, blue), and from 2D poses obtained by projecting ground truth 3D poses in the ventral x-y plane (*α*′, *β*′, *γ*′, *ω*′, **Figure S1**, green). Due to the uncertainty of 3D pose estimation, we aimed to provide upper and lower confidence bounds. Therefore, we assumed that the keypoint coordinates would be Gaussian distributed around the estimated 3D coordinate. As a proxy for the variance we took the variation of bone lengths because they are expected to remain approximately constant owing to the low mechanical compliance of the fly’s exoskeleton (with the exception of the flexible tarsal segments). This allowed us to predict 3D joint angles by Monte Carlo sampling (see Materials and Methods).

We found that joint angles derived from lifted and triangulated 3D poses were in close agreement (**Figure S1**, red and blue). The errors are also low when comparing angle estimate variances to the amount of joint rotation during locomotor cycles. This shows that our network learned and preserved body proportions—a remarkable fact given the absence of any skeletal constraints, or temporal information. Furthermore, when comparing the joint angles derived from 3D and 2D poses, we found that the predicted coxa-femur 3D joint angles, *β*, in the front and hindlegs were of larger amplitude than their projected 2D counterparts, *β*′. This is expected since the action of these joints has a large out-of-plane component relative to the projected x-y plane during walking. Second, in the front leg, the predicted tibia-tarsus 3D joint angles, *ω*, were of smaller amplitude than their projected 2D counterparts, *ω*′. Indeed, rotations upstream in the kinematic chain (proximal joints) cause the movement of the whole leg, which can introduce spurious variations in the angles of distal joints when viewed from a projected plane. These results illustrate how 3D poses predicted by LiftPose3D can help to decouple the underlying physical degrees-of-freedom and avoid spurious correlations introduced by 2D projected joint angles.

Because LiftPose3D maintained prediction accuracy irrespective of viewing angle (**Figure 1F**), we next asked how it would perform when predicting 3D poses in freely behaving animals, where the effective camera angle dynamically changes. We were also interested in considering animals without exoskeletons where nearby keypoint movements are less constrained. We addressed this question by training LiftPose3D to predict 3D poses for freely behaving macaque monkeys recorded in the OpenMonkeyStudio dataset [8]. These data consist of 3D poses obtained by triangulating markerless 2D pose estimates [42] from 62 calibrated, synchronized, and distributed cameras (**Figure 1H**). After training the network with only 6’571 3D poses, we could lift 3D poses from test images— including macaques walking as well as taking up diverse poses (**Video 4**)—from any of the 62 cameras (**Figure 1I**), and with a relatively small body length-normalized MAE (**Figure 1J**).

Taken together, these results demonstrate that LiftPose3D can reduce the number of cameras required to perform full and accurate 3D pose estimation with simple data preprocessing and a relatively small but diverse training dataset.

### 2.3 Predicting 3D pose with occluded keypoints in freely behaving animals

In freely behaving animals, keypoints are often missing from certain camera angles due to self-occlusions and, therefore, only partial ground truth 3D annotations can be obtained by triangulation. We asked how the global nature of lifting—all keypoints are lifted simultaneously—might be leveraged to reconstruct information lost by occlusions and to predict full 3D poses.

To address this question, we built an experimental system consisting of a transparent enclosure physically coupled to a right-angle prism mirror, similar to previous recording systems used for flies and mice [18, 43, 44]. We used a single camera beneath the platform to record the ventral and side views of a freely behaving fly (**Figure 2A**) and trained two DeepLabCut models [6] to obtain 2D joint coordinates from each of these views (**Figure 2A**). Having only two views meant that keypoints closer to the prism were simultaneously visible in both views and could therefore be triangulated, while those occluded from the side view had only ventral 2D information, which is insufficient for triangulation. With this partial 3D ground truth, it was thus *a priori* unclear if a LiftPose3D network could be trained to lift 3D poses using only ventral 2D poses (**Figure 2A**, green box).

**Figure 2:**
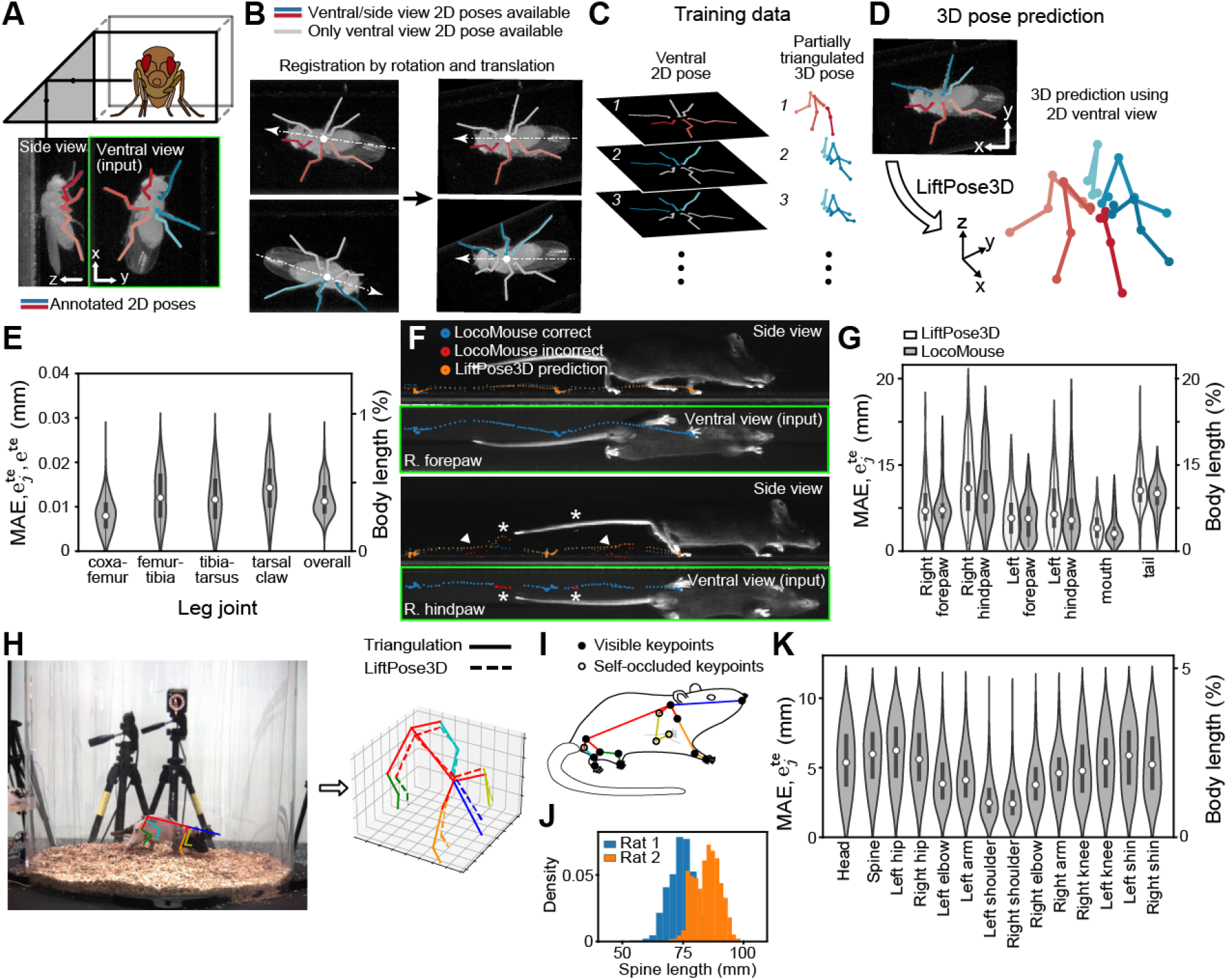
LiftPose3D performs 3D pose estimation on freely behaving animals with occluded keypoints. **A** *Drosophila* behaving freely within a narrow, transparent enclosure. Using one camera and a right-angle prism mirror, both ventral (top) and side (bottom) views are visible. 2D poses are tracked using two separately trained deep networks for each view (colored lines). Ventral 2D poses (green box) are used for lifting the 3D pose. **B** Keypoints near the prism mirror (red and blue) can be tracked in both views and triangulated. The remaining keypoints (gray) are only visible in the ventral view and thus have no 3D triangulated ground truth. To obtain triangulated ground truth examples for both sides of the bilaterally symmetric fly, we register the ventral images to align the orientation and position of all animals. **C** Training data thus consists of a set of full ventral view 2D poses and their corresponding partially triangulated 3D poses. **D** Following training with these aligned 2D-3D ground truth poses, LiftPose3D can be used to predict 3D poses for new ventral view 2D pose data. **E** Joint-wise and overall absolute errors of the network’s 3D pose predictions for freely behaving *Drosophila*. **F** A similar data preprocessing approach can be used to lift ventral view 2D poses of mice (green boxes) walking within a narrow enclosure and tracked using the LocoMouse software. LocoMouse ground truth (blue and red) and LiftPose3D (orange) pose trajectories are shown for the right forepaw (top) and hindpaw (bottom) for one walking epoch. Arrowheads indicate where LiftPose3D lifting of the ventral view can be used to correct LocoMouse side view tracking errors (red). Asterisks indicate where inaccuracies in the LocoMouse ventral view ground truth (red) disrupt LiftPose3D’s side view predictions (orange). **G** Absolute errors of LiftPose3D and LocoMouse side view predictions for six keypoints with respect to a manually-annotated ground truth. **H** LiftPose3D can be trained to lift 3D poses of a freely moving rat with occluded keypoints. **I** Large animal-to-animal skeleton variation illustrated by histograms of the measured lengths of the spinal segment for two animals. **J** Camera image from the CAPTURE dataset superimposed with the annotated 2D pose (left). LiftPose3D uses this 2D pose to recover the full 3D pose (right). **K** Error distribution over all keypoints for the CAPTURE dataset.

Since the ventral and side views enclose right angles (i.e., are orthographic projections of the true 3D pose), and because long focal length cameras have negligible perspective effects, we used 2D poses from the ventral view to estimate the *z*-axis depth of occluded keypoints in the unseen side view. Because all keypoints were simultaneously visible from the ventral view, this allowed us to align flies in the same reference frame (**Figure 2B**), and transform lifting to the regression problem in Eq. (1) where the indicator function 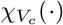 now represents the visible keypoints from the side camera (**Figure 2C**). As a result, keypoints with incomplete 3D information were not penalized during training. Taking the ventral view as an input, where we all keypoints were present, but penalizing only those with complete 3D information allowed the network to implicitly regress the unseen coordinates during training. We found that LiftPose3D could also predict 3D positions for every joint at test time, including those occluded in the prism’s side view (**Figure 2D** and **Video 5**). Notably, the accuracy, based on available triangulation-derived 3D positions (**Figure 2E**) was better than that obtained for tethered flies by triangulation with four cameras (**Figure 1E**). Thus, LiftPose3D can estimate 3D poses from 2D images in cases where keypoints are occluded and cannot be triangulated.

These results suggested an opportunity to apply lifting to identify and potentially correct in-accurate 3D poses obtained with other approaches. We considered a previously published dataset consisting of freely behaving mice traversing a narrow corridor [18] tracked by the LocoMouse software from both ventral and side views [18]. Using these, we triangulated incomplete 3D ground truth poses (due to side view occlusions) and, as in the *Drosophila* prism mirror dataset, placed them in the same reference frame by registering the ventral poses. We then trained a LiftPose3D network to lift the ventral 2D poses (**Figure 2F**, green boxes). Predictions were in good agreement with the LocoMouse’s side view tracking (**Figure 2E** and **Video 6**) and could recover expected cycloid-like kinematics between strides (**Figure 2F**). Remarkably, LiftPose3D predictions could also correct side-view poses that were incorrectly labeled or missing in the ground truth dataset (**Figure 2F**, bottom, white arrowheads). However, lifting accuracy depended on the fidelity of input 2D poses: incorrect ventral 2D poses generated false side view predictions (**Figure 2F**, bottom, white asterisks). These errors were always restricted to the joint-of-interest and were relatively infrequent. Overall, LiftPose3D performed as well as LocoMouse, when compared with manual human annotation (**Figure 2G**). These results demonstrate that LiftPose3D can be used to correct other tracking methods, but also highlights the importance of quantifying the confidence of input 2D poses to avoid lifting keypoints incorrectly.

The above examples demonstrate that LiftPose3D learns spatial relationships between keypoints when they are presented in the same reference frame. We therefore asked how well this feature generalizes to animals generating more complex behaviors and with large variations in body proportions. As an example, we considered a recently published CAPTURE dataset that used six fixed cameras to record freely moving rats within a circular naturalistic arena [39] (**Figure 2H, left**). The keypoints were visual markers placed on the fur of the animals. These were intermittently self-occluded during motion (**Figure 2I**). Moreover, these animals performed a variety of complex behaviors including walking, reaching, rearing, and turning. During these movements, 2D pose skeletons underwent large deformations. This is illustrated by the broad distribution of keypoint distances conveying spine lengths (**Figure 2J**). Despite these challenges we aimed to train a lifting network for these data, thus requiring a series of further innovations. First, to overcome the variations in body proportions both within and across animals, we first constructed a template skeleton with bone lengths that followed independent normal distributions with means and standard deviations representative of expected bone lengths across the population of recorded animals. During training, we randomly sampled from these distributions to rescale each ground truth 3D pose while preserving joint angles. Then, we obtained a corresponding 2D pose via projection. Second, although the animal-to-camera angle changed continuously during animal behaviors, we augmented the training data by generating virtual 2D projections within the Euler angle range of ±10° about all three axes. Third, although the depth-wise motion of animals caused substantial variation in their distance to the camera, we assumed that it remained large enough for the weak perspective condition to hold, and normalized 2D poses by their Frobenius norm, as before (see Materials and Methods). By doing so, the camera parameters at test time no longer needed to be known, making our network directly applicable to other rat movement studies. To illustrate this, we trained our network on two experiments from the CAPTURE data (consisting of two animals and two camera arrangements) and then tested it on a third experiment with a different animal and camera arrangement (i.e., different focal length and orientation). In each case, we presented zeros to the network in place of missing data points and found that LiftPose3D could accurately predict the nonzero coordinates (**Figure 2H, right and K, Video 7**). This shows that erroneous 2D point coordinates, which would otherwise confound lifting performance (**Figure 2F**), can be dealt with by presenting zeros in place of low confidence points. Additionally, our methods could largely compensate for the challenges associated with lifting 3D poses for freely behaving animals having large variations in body proportions.

### 2.4 Using domain adaptation to lift diverse experimental data when triangulation is impossible

Our angle-invariant lifter networks for tethered flies (**Figure 1D-F**) and for freely behaving rats (**Figure 2**H-K) can be directly used in similar experimental systems without having to collect additional 3D pose training data. However, small variations in new experimental systems resulting from camera distortion or postural differences may limit the accuracy of lifted 3D poses. Therefore, the possibility of domain adaptation–using pretrained networks to lift poses in new experimental scenarios with small postural variations–could enable extending the value of LiftPose3D to a vast and diverse user group who have only a single-camera for acquiring 2D poses and no means to obtain a ground truth library of 3D poses.

We assessed the possibility of domain adaptation by training a network in domain *A*—tethered flies on a spherical treadmill—and predicting 3D poses in domain *B* —freely-moving flies on a flat surface (**Figure 3A**). We chose this pair of experiments due to the availability of ground truth data in both domains, which we could use to measure accuracy. Before performing domain adaptation, we first derived poses from 2D ventral images in domain *B*, as before. This allowed us to circumvent the difficulties arising from differences in appearance and illumination that are present in the more general image domain adaptation problem [45, 46]. Thus, adapting poses became a purely geometric problem of adjusting proportions and postural differences across domains. **Figure 3A** depicts the three-step process to lift a 2D pose in domain *B*. First, we used a linear transformation *d*_2_ to transform the 2D pose into the source domain *A*. Second, we lifted this 2D pose into a 3D pose using a LiftPose3D network pre-trained only on 3D poses from domain *A*. Third, we transformed the lifted 3D poses from domain *A* back to domain *B* using another linear transformation *d*_3_. To find *d*_2_ and *d*_3_, we identified, for every pose in a training dataset *B’, k* nearest neighbors *aï* in domain *A* (**Figure 3A,B**), and used these to find the best-fit linear transformations between domains (see Materials and Methods for details). These linear transformations are expected to generalize as long as the poses in domain *A* are rich enough to cover the pose repertoire in domain *B* and are sufficiently similar between domains. We tested this by 10-fold cross-validation (with *k* = 1 for *d*_2_ and *k* = 2 for *d*_3_) and found that the error associated with the transformations converged after less than 500 poses (**Figure 3C**). The final lifted poses were also in good agreement with the triangulated poses in domain *B* (**Figure 3D**). The accuracy was slightly worse but remarkably comparable with that of a network lifting purely in domain *A* (**Figure 3E, compare dark with light gray**).

**Figure 3:**
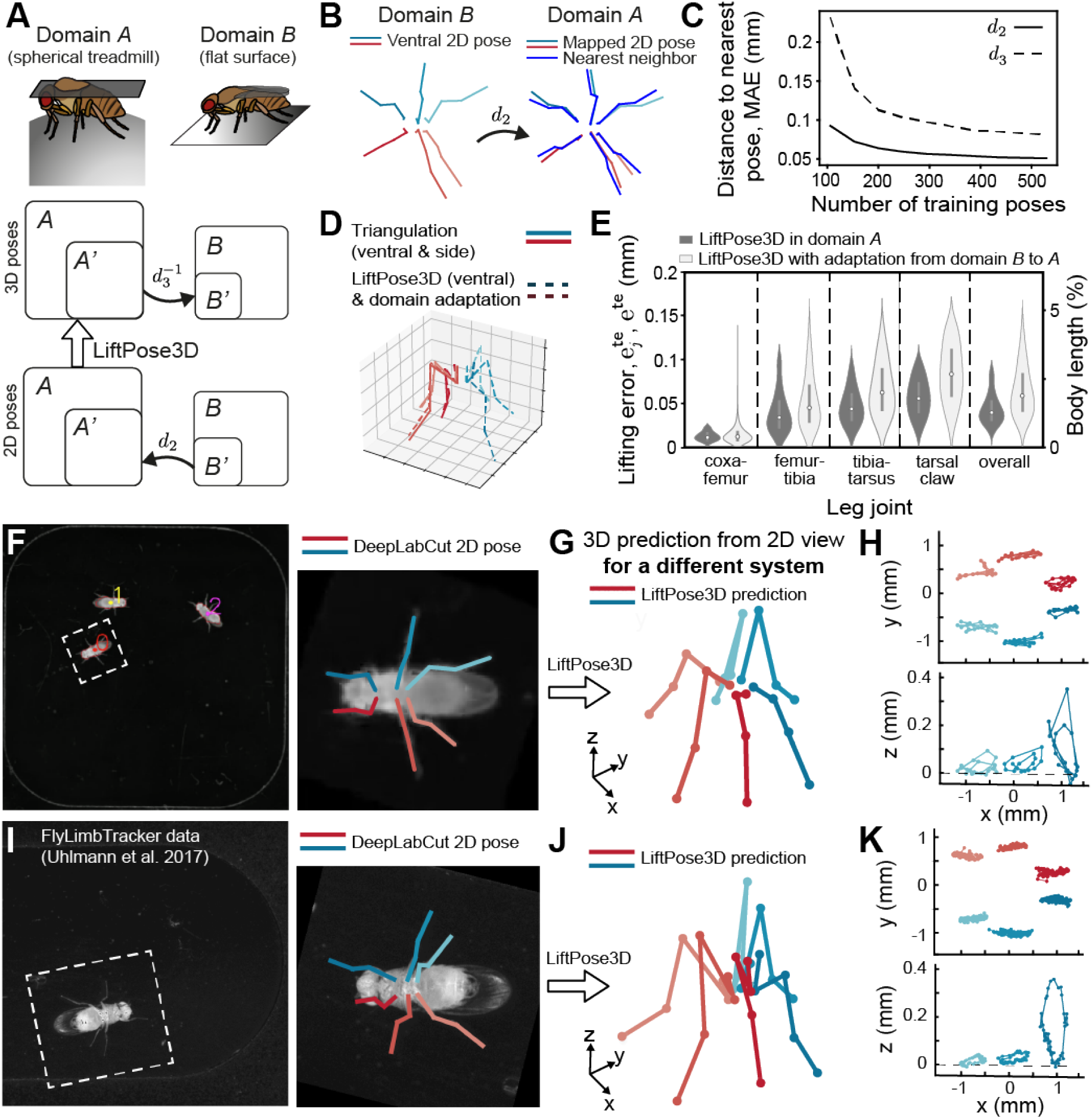
A pretrained LiftPose3D network predicts 3D poses for diverse data and when triangulation is impossible. **A** Linear domain adaptation between domain A (fly on a spherical treadmill) and domain B (fly on a flat surface). 2D poses in B are mapped to A by a linear transformation *d*_2_ then lifted with a network trained only on domain A poses. After lifting, the 3D poses are mapped back to B by another linear transformation *d*_3_. **B** A typical 2D pose in domain B mapped into domain A by the best-fit linear transformation *d*_2_ between poses in B and their nearest neighbors in A. **C** Error between mapped pose and nearest neighbor poses for *d*_2_, *d*_3_ against the number of poses used to train them. The number of nearest neighbors used was *k* = 1 fof *d*_2_ and *k* = 2 for *d*_3_. **D** Lifted 3D pose following domain adapation of a ventral domain B 2D pose and lifting with a network trained on domain A data. The prediction is superimposed with the imcomplete ground truth 3D pose in domain B. **E** Lifting error following domain adaptation of domain B poses compared with lifting error in the domain A with no domain adaptation. **F** Freely behaving flies recorded from below using a low-resolution camera. Following body tracking, the region-of-interest containing the fly is cropped and registered. 2D pose estimation is then performed for the 24 visible joints. **G** 2D poses are adapted to the prism-mirror domain. These are then lifted to 3D poses with pre-trained network using prism-mirror data and coarse-grained to match the lower resolution 2D images in the new experimental system. **H** These 3D poses permit the analysis of claw movements in the otherwise unobserved *x − z* plane (bottom). **I** Published data from [20] showing a freely behaving fly recorded from below using one high-resolution camera. 2D pose estimation was performed for all 30 joints. Following tracking, a region-of-interest containing the fly was cropped and registered. The same LiftPose3D network trained in panel B—but without coarse-graining—was used to predict **J** 3D poses and **K** unobserved claw movements in the *x − z* plane (bottom).

To demonstrate the full potential of domain adaptation, we next focused on lifting *Drosophila* 2D poses recorded from a single ventral camera. This approach is the most widely used free behavior paradigm in laboratory settings due to its simplicity, low-cost, and increased throughput. It has been applied to study many organisms including *C. elegans* [47], larval zebrafish [48], larval *Drosophila* [49], adult *Drosophila* [50], and mice [51]. Although these recordings can be augmented with depth sensors [52,53], such sensors cannot resolve small laboratory animals, or reconstruct full 3D poses. Thus, 3D pose estimation of laboratory animals from a single 2D view remains an unsolved and highly desirable goal, with the potential to substantially enrich behavioral datasets and to improve downstream analysis.

First, we developed a new experimental system consisting of a square-shaped arena in which multiple freely-behaving flies could be recorded ventrally using a single camera (**Figure 3F**, left). In addition to being a different experimental system from our prism mirror setup and using a different camera, here the images had four-fold lower spatial resolution (26 px mm^*−*1^). Hence, we could only label 24 visible keypoints using DeepLabCut (**Figure 3F**, right). We then pretrained a network using prism-mirror training data—using only the keypoints present in both datasets—and augmented these data using a Gaussian noise term with standard deviation of ∼4 (see Materials and Methods). Before lifting, we domain-adapted the annotated 2D poses into the network’s domain, as before (**Figure 3B**). Because ventrally-viewed leg configurations during swing and stance phases are difficult to distinguish, particularly at lower resolution, to reconstruct realistic joint movements our network would have to first learn the postural relationships between each leg. Remarkably, we found that the network could predict physiologically realistic 3D poses in this new dataset using only ventral 2D poses (**Figure 3G** and **Video 8**). During walking, 2D tracking of the tarsal claws traced out stereotypical trajectories in the x-y plane (**Figure 3H**, top) [54] and circular movements in the unmeasured x-z plane (**Figure 3H**, bottom) whose amplitudes were consistent with real kinematic measurements during forward walking [55].

The ability to adapt training data from one domain to another also raises the exciting possibility that LiftPose3D could be used to ‘resurrect’ previously published 2D pose data for new 3D kinematic analysis. To test this, we applied our prism mirror-based training data to lift previously published high-resolution (203 px mm^*−*1^) video data of a fly walking freely through a capsule-shaped arena [20] (**Figure 3I**). Using a similar data processing pipeline as for the previous case (**Figure 3B,F,G**), including registration and domain adaptation but not noise perturbations (the target data were of similarly high resolution as the training data), the LiftPose3D network could effectively predict 3D poses from this previously published dataset (**Figure 3J**). We again observed physiologically realistic cyclical movements of the pretarsi during forward walking (**Figure 3K**, bottom; **Video 9**). Thus, thanks to the adaptation of pretrained networks to new domains, LiftPose3D can be an effective tool for performing 3D pose estimation on previously published 2D video data for which 3D triangulation would be otherwise impossible.

### 2.5 *Drosophila* LiftPose3D station

These domain adaptation results opened up the possibility to make 3D pose acquisition considerably cheaper and more accessible across laboratories. To explore this possibility, we developed and constructed a “*Drosophila* LiftPose3D station” consisting of an inexpensive (∼$150) open-source hardware system including a 3D printed rig supporting a rectangular arena recorded by a Raspberry Pi camera and illuminated using LEDs (see **Figure S3** and Materials and Methods). A common hardware solution like this one eliminates compounding variables introduced across different experimental setups (e.g., camera distortion and perspective effects) and allowed us to provide pre-trained DeepLabCut and LiftPose3D networks that permit straightforward 3D pose measurements by other laboratories for *Drosophila* behavioral studies (**Video 10**). We envision that such an approach— a common behavioral arena, camera and illumination hardware, and pretrained pose estimation networks—might, in the future, also facilitate cross-laboratory lifting of mouse 2D poses using a single camera.

## 3 Discussion

Here we have introduced LiftPose3D, a deep neural network-based tool that dramatically simplifies and enables 3D pose estimation for a wide variety of laboratory contexts. Our approach uses the network architecture of [34], originally designed for human-pose estimation, and introduces a series of innovations to input data preprocessing, training augmentation and domain adaptation. These contributions enable network training with several orders of magnitude less training data and when ground truth 3D poses are incomplete due to occlusions or corrupted by inaccurate labelling. We have also developed data augmentation methods that make LiftPose3D networks invariant to camera hardware and positioning, allowing them to generalize across arbitrary setups. Furthermore, we provide a comprehensive software pipeline for data preprocessing, network training, 3D predictions, and visualization. A single intuitive Python notebook interfaces all the tools needed to obtain the results shown here.

We illustrate how LiftPose3D reduces the number of cameras required for 3D pose estimation; from three to one on each side of a tethered fly, and from 62 to one in freely behaving macaques. In the case of flies, we also describe the training of a camera hardware-invariant network that can take inputs from any low-distortion camera positioned at an arbitrary orientation relative to the target animal. We also provide two pre-trained networks—one for a side-view camera placed at any orientation and one for a ventral camera—that can be readily used for new experimental systems. In all cases, high accuracy comparable to triangulation was achieved for a range of both stereotypic and irregular spontaneous behaviors. For freely behaving flies, mice and rats, we have demonstrated that LiftPose3D can estimate 3D poses despite self-occlusions and that it can identify and correct keypoints that have been mislabeled by other keypoint tracking approaches. Finally, we have demonstrated that linear domain adaptation can be used to account for variations due to camera distortion or animal poses in new datasets. We used this approach to predict 3D poses for flies moving freely on a flat surface with a LiftPose3D network pre-trained with data of tethered flies on a spherical treadmill. Domain adaptation also opens up the possibility to acquire 3D pose data in situations where 3D ground truth is impossible to obtain by multi-camera triangulation, including lifting 3D poses from a large corpus of previously published 2D video data for further kinematic analysis. Using our domain adaptation methodology, networks with the largest and most diverse training data, like that for the tethered fly—may already be sufficiently robust to accurately lift 2D to 3D pose in other laboratories. To capitalize on this, we developed and demonstrate how this can be applied with an inexpensive open hardware platform, the LiftPose3D station. Setups like this will dramatically lower the barrier for 3D pose estimation in other laboratories around the world.

The LiftPose3D framework is general and can be applied with very few changes to study different laboratory animals in new experimental systems and with diverse data acquisition rates, image resolutions, and 2D pose input sources including—as we demonstrate in this study—the stacked hourglass network of DeepFly3D [7] and DeepLabCut [6]. Nevertheless, several factors must be taken into consideration when optimizing LiftPose3D for new experimental systems. First, because predicting depth from a 2D projection depends on comparing the projected lengths of body parts, input poses must be sufficiently well-resolved to discriminate between 3D poses that have similar 2D projections. Second, prediction accuracy depends on the diversity of training data—i.e., measured behaviors. We caution that previously untrained behaviors may not be as accurately lifted using a pretrained network. In the future, we envision that robust lifting networks might be generated by a communal, inter-laboratory aggregation of 3D pose ground truth datasets that include a variety of spontaneously generated and experimentally-induced behaviors. Third, although our aim was to develop a general tool with minimal experiment or animal-specific features, further work can improve LiftPose3D predictions for specific applications by bootstrapping to 3D body priors, thereby constraining the space of possible 3D poses [56–60]. Finally, lifting might also be improved by using a network that incorporates temporal information for data acquired at a constant frame rate [35].

We anticipate that LiftPose3D can already accelerate the successful adoption of 3D pose estimation in laboratory research by reducing the need for complex and expensive synchronized multi-camera systems, and arduous calibration procedures. This, in turn, will improve the fidelity and quality of behavioral kinematic data needed to understand how actions emerge from multi-scale biological processes ranging from gene expression to neural dynamics and biomechanics.

## 4 Materials and Methods

### 4.1 Obtaining 3D pose ground truth data by triangulation

To obtain the 3D ground truth coordinates **X**_*j*_ ∈ ℝ^3^ for joints *j* = 1, …, *n* from a set of 2D keypoints **x**_*c,j*_ ∈ R^2^ in images acquired by the cameras *c* = 1, …, *N* we followed the procedure described in [7].Let us express 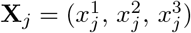 in homogeneous coordinates as 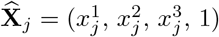. The projection from the 3D points in the global coordinate system to 2D points in a local coordinate system centered on camera *c* is performed by the function *π*_*c*_ : ℝ^4^ →ℝ^3^ defined as 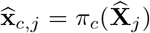. This function can be expressed as a composition *π*_*c*_ = proj_1,2_ *ϕ*_*c*_ of an affine transformation *ϕ*_*c*_ : ℝ^4^ →ℝ^4^ from global coordinates to camera-centered coordinates and a projection proj_1,2_ : ℝ^4^ →ℝ^3^ to the first two coordinates. Both functions can be parametrized using the pinhole camera model [14]. On the one hand, we have

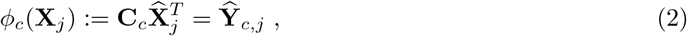

where **C**_*c*_ is the extrinsic camera matrix corresponding to the *ϕ*_*c*_ and can be written as

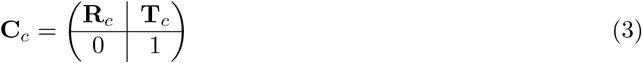

where **R**_*c*_ ∈ ℝ^3*×*3^ is a matrix corresponding to rotation around the origin and **T**_*c*_ ∈ ℝ^3^ is a translation vector representing the distance of the origin of the world coordinate system and the camera center. Likewise, the projection function can be expressed as

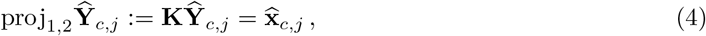

where **K** is the intrinsic camera transformation

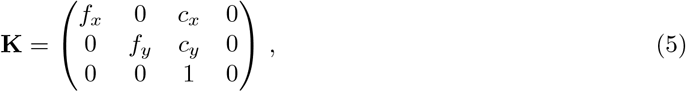

where *f*_*x*_, *f*_*y*_ denote the focal lengths and *c*_*x*_, *c*_*y*_ denote the image center. The coordinates projected to the camera plane can be obtained by converting back to Euclidean coordinates 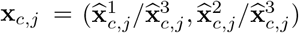.

Triangulation of the coordinate **X**_*j*_ of joint *j* with respect to *π*_*c*_ is obtained by minimizing the reprojection error, that is, the discrepancy between the 2D camera coordinate, **x**_*c,j*_, and the 3D coordinate projected to the camera frame, *π*_*c*_(**X**_*j*_). Let *V*_*c*_ be the set of visible joints from camera *c*. The reprojection error for joint *j* is taken to be

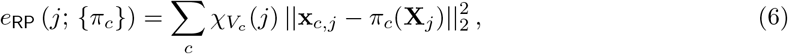

where 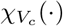 is the indicator function of set *V*_*c*_ of visible keypoints from camera *c*. The camera projection functions *π*_*c*_ are initially unknown. To avoid having to use a calibtration grid, we jointly minimize with respect to the 3D location of all joints and to the camera parameters, a procedure known as bundle adjustment [14]. Given a set of 2D observations, we seek

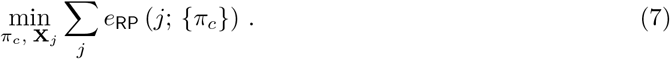

using a second-order optimization method. For further details, we refer the interested reader to [7].

### 4.2 LiftPose3D network architecture and optimization

The core LiftPose3D network architecture is similar to the one of [34] and is depicted by **Figure 1B**. Its main module includes two linear layers of dimension 1024 rectified linear units (ReLU, [61]), dropout [40] and residual connections [62]. The inputs and outputs of each block are connected during each forward pass using a skip connection. The model contains 4 ×10^6^ trainable parameters, which are optimized by stochastic gradient descent using the Adam optimizer [63]. We also perform batch normalization [64].

In all cases, the parameters were set using Kaiming initialization [62] and the optimizer was run until convergence—typically within 30 epochs—with the following training hyperparameters: Batch-size of 64 and an initial learning rate of 10^*−*3^ that was dropped by 4% every 5000 steps. We implemented our network in PyTorch on a desktop workstation running on an Intel Core i9-7900X CPU with 32 GB of DDR4 RAM, and a GeForce RTX 2080 Ti Dual O11G GPU. Training time was less than 10 minutes for all cases studied.

### 4.3 Camera-angle augmentation

The object-to-camera orientation is encoded by the extrinsic matrix **C**_*c*_ of Eq. 3. When it is unavailable, one can still use our framework by taking 3D poses from the ground truth library and, during training, performing virtual 2D projections around the approximate camera location or for all possible angles. To this end, we assume that the rotation matrix **R** is unknown, but that the intrinsic matrix **K** and the object-to-camera distance *d* are known such that we may take **T** = (0, 0, *d*)^*T*^. When **K** or *d* are also unknown, or dynamically changing, one can make the weak-perspective assumption as in described in the next section. Then, instead of training the LiftPose3D network with pairs of 3D poses and 2D poses at fixed angles, we perform random 2D projections of the 3D pose to obtain virtual camera planes whose centers *c*_*x*_, *c*_*y*_ lie on the sphere of radius *d*. To define the projections we require a parametric representation of the rotations. Rotating a point in 3D space can be achieved using three consecutive rotations around the three Cartesian coordinate axes *x, y, z* commonly referred to as Euler angles and denoted by *ψ*_*x*_,*ψ*_*y*_, and *ψ*_*y*_. The rotation matrix can then be written as

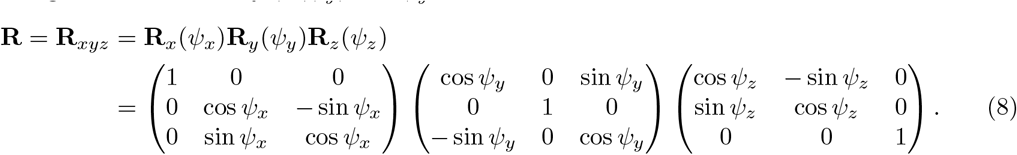

Given Eq. (2)–(5) we may then define a random projection 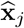 on the sphere of radius *d* of a keypoint with homogeneous coordinate 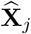 as

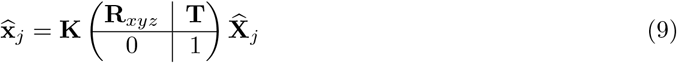

where **T** = (0, 0, *d*)^*T*^. Likewise, the 3D pose in camera coordinates can be expressed as

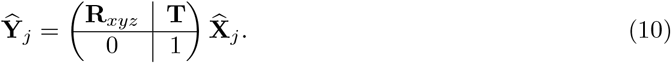

Before training, we fix *d, f*_*x*_, *f*_*y*_, *c*_*y*_, *c*_*y*_ and define intervals for the Euler angle rotations. We then obtain the mean and standard deviation in each dimension both for 2D and 3D poses in the training data set by performing random projections within these angle ranges. The obtained means and standard deviations are used to normalize both the training and test datasets.

### 4.4 Weak perspective augmentation

To project 2D pose from 3D pose, one needs to know the camera transformation *ϕ*_*c*_ (Eq. (2)), encoded by the extrinsic matix **C**_*c*_ (Eq. (3)) and the projection function proj_1,2_ (Eq. (4)), encoded by the intrinsic matrix **K** (Eq. (5)). In the previous section, we described how to deal with the case when **C**_*c*_ is unknown. In addition, **K** may also be unknown *a priori* at test time. Alternatively, one may want to use one of our pre-trained networks on a novel dataset without having to match the camera positioning (focal length, camera-to-animal distance) used to collect the training data. In this case, one may still be able to predict the 3D pose in a fixed camera-centered coordinate frame by assuming that either the camera-to-animal distance or the focal length are large enough to neglect perspective effects and by normalizing the scale of 2D poses. Following Ref. [41], we choose the Frobenius norm to perform normalization on the input 2D poses ||**x**_*c,j*_*/* **x**_*c,j*_|| _*F*_, which is the diagonal distance of the smallest bounding box around the 2D pose. Note, that if the 2D poses are obtained via projections, one may use the unit intrinsic matrix Eq. (5) with *f*_*x*_ = *f*_*y*_ and *c*_*x*_ = *c*_*y*_ = 0 before performing normalization. Here, using *c*_*x*_ = *c*_*y*_ = 0 assumes that the 2D poses are centered, which in each of our examples is achieved by considering coordinates relative to root joints placed at the origin. Importantly, the 2D poses must be normalized both during training and test times.

### 4.5 Linear domain adaptation

Here we describe the process of adapting a network trained on data from experiment *A* to lift 2D poses in experiment *B*. Domain adaptation is also useful if the camera parameters or the distance from the camera are not known and the weak perspective assumption cannot be invoked.

Here, the basis for domain adaptation is to first find a function *d*_2_ : *B*|_2_ → *A*|_2_, where *A*|_2_ and *B*|_2_ are restrictions of 3D poses in the two domains to the corresponding 2*n*-dimensional spaces of 2D poses. This function maps poses in domain *B* to domain *A* and makes them compatible inputs for the network trained on poses in domain *A*. In the scenario that 3D data is available in domain *B*, we can also find a function *d*_3_ : *B* →*A* where *A* and *B* are 3*n*-dimensional spaces of 3D poses in the two experimental domains. After 3D poses have been obtained in domain *A*, we map back these poses to domain *B* by inverting this function.

We now describe how to obtain the functions *d*_2_ and *d*_3_, which we denote collectively as *d*. To find *d*, we assume that poses in domain *B* can be obtained by small perturbations of poses in domain *A*. This allows us to set up a matching between the two domains by finding nearest neighbor 2D poses in domain *A* for each 2D pose in domain *B*, 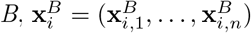. We use 2D rather than 3D poses to find a match because 3D poses may not always be available in domain *B*. Moreover, the nearest poses in 3D space will necessarily be among the nearest poses in 2D space. Specifically, for each 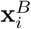, we find a set of *k* nearest poses in domain 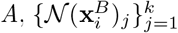 such that 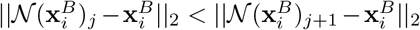. We then use these poses to learn a linear mapping **W**_*BA*_ ∈ ℝ^2*n×*2*n*^ from domain *B* to *A*, where *n* is the number of keypoints, as before. We can find this linear mapping by first defining a set of *p* training poses in domain 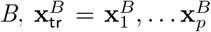 and writing 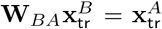, where 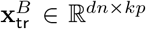 and 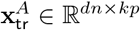 with *d* = 2 or 3 are matrices defined according to

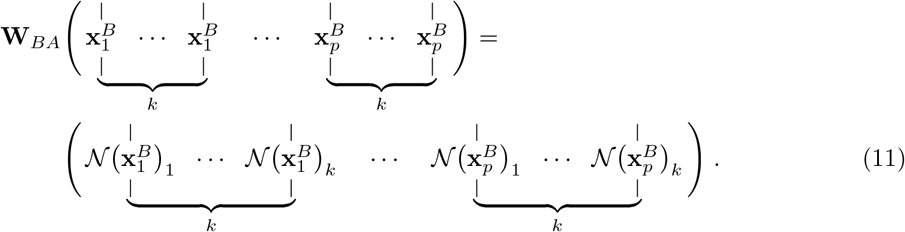

Transposing this linear equation yields the linear problem 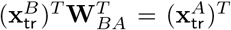. Given that the *p* training poses are different, 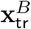 has linearly independent columns and this problem is overdetermined as long as *kp > dn*. Thus, by least-squares minimization, we obtain 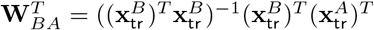.

### 4.6 Experimental systems and conditions

All adult *Drosophila melanogaster* experiments were performed on female flies raised at 25°C on a 12 h light/dark cycle at 2-3 days post-eclosion (dpe). Before each experiment, wild-type (*PR*) animals were anaesthetized using CO_2_ or in ice-cooled vials and left to acclimate for 10 min. DeepFly3D tethered fly data were taken from [7]. OpenMonkeyStudio macaque data were taken from [8]. LocoMouse mouse data were taken from [18]. CAPTURE rat data were taken from [39]. FlyLimbTracker freely-behaving fly data were taken from [20]. See these publications for detailed experimental procedures. For more information on the datasets including the number of keypoints, poses, animals, resolution, framerate we refer the reader to **Table 1**.

**Table 1:**
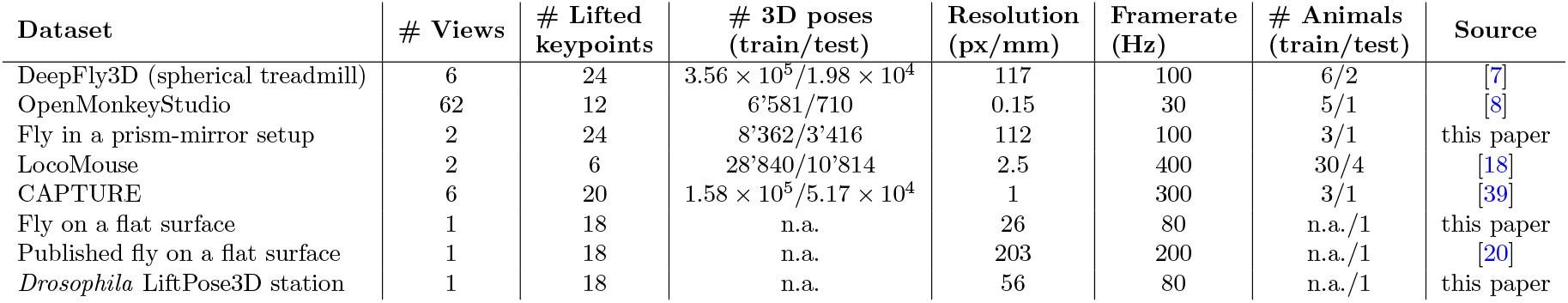
List of datasets used

#### 4.6.1 Freely behaving *Drosophila* recorded from two high-resolution views using one camera and a right-angle prism mirror

We constructed a transparent arena coupled to a right-angle prism mirror [43,44]. The enclosed arena consists of three vertically stacked layers of 1/16” thick acrylic sheets laser-cut to be 15 mm long, 3 mm wide, and 1.6 mm high. The arena ceiling and walls were coated with Sigmacote (Sigma-Aldrich, Merck, Darmstadt, Germany) to discourage animals from climbing onto the walls and ceilings. One side of the enclosure was physically coupled to a right-angled prism (Thorlabs PS915). The arena and prism were placed on a kinematic mounting platform (Thorlabs KM100B/M), permitting their 3D adjustment with respect to a camera (Basler acA1920-150um) outfitted with a lens (Computar MLM3X-MP, Cary, NC USA). The camera was oriented vertically upwards below the arena to provide two views of the fly: a direct ventral view, and an indirect, prism mirror-reflected side view. The arena was illuminated by four Infrared LEDs (Thorlabs, fibre-coupled LED M850F2 with driver LEDD1B T-Cube and collimator F810SMA-780): two from above and two from below. To elicit locomotor activity, the platform was acoustically and mechanically stimulated using a mobile phone speaker. Flies were then allowed to behave freely, without optogenetic stimulation.

#### 4.6.2 Freely behaving *Drosophila* recorded from one ventral view at low-resolution

We constructed a square arena consisting of three vertically stacked layers of 1/16” thick acrylic sheets laser-cut to be 30 mm long, 30 mm wide, and 1.6 mm high. This arena can house multiple flies at once, increasing throughput at the expense of spatial resolution (26 px mm^*−*1^). Before each experiment the arena ceiling was coated with 10 uL Sigmacote (Sigma-Aldrich, Merck, Darmstadt, Germany) to discourage animals from climbing onto the ceiling. A camera (pco.panda 4.2 M-USB-PCO, Gloor Instruments, Switzerland, with a Milvus 2/100M ZF.2 lens, Zeiss, Switzerland) was oriented with respect to a 45 degree mirror below the arena to capture a ventral view of the fly. An 850 nm infrared LED ring light (CCS Inc. LDR2-74IR2-850-LA) was placed above the arena to provide illumination. Although the experiment contained optogenetically elicited behaviors interspersed with periods of spontaneous behavior, here we focused only on spontaneously generated forward walking.

The positions and orientations of individual flies were tracked using custom software including a modified version of Tracktor [65]. Using these data, a 138 ×138 px image was cropped around each fly and registered for subsequent analyses.

#### 4.6.3 *Drosophila* LiftPose3D station

The LiftPose3D station is an easily constructed and used system designed to capture 2D poses of freely behaving *Drosophila melanogaster*. The station is powered by a Raspberry Pi Zero board and uses a high quality camera with a 6 mm wide-angle lens to obtain images at 800×800 pixel resolution. The cameraïs exposure time was set to 2 ms and its framerate to 80 fps. Images are first stored as jpeg files in the micro SD card of the Raspberry Pi Zero, and then transfered to a workstation for further processing. Each image file size is about 25 kb. Therefore we are able to store up to 3 hrs of data using our current configuration. We refer the reader to the Supplementary Notes for a detailed description of the design and assembly. **Table 2** provides a full list of components with links to retailers from whom they can be purchased, or computer-aided designs (CAD) of custom manufactured pieces.

**Table 2:** List of components composing the LiftPose3D station

| Quantity | Component | Company/<br>Manufacturing method | Type (Alternative)/<br>Material (CAD) |
| --- | --- | --- | --- |
| 1 | Raspberry Pi | Raspberry Pi | Zero W |
| 1 | Raspberry Pi Camera | Raspberry Pi | High Quality Camera |
| 1 | Raspberry Pi CS-mount lens | CGL | 6 mm wide-angle |
| 1 | Raspberry Pi Zero Flex Cable | Sertronics | RPIZ-FLEX-15 |
| 1 | Micro SD memory | Sandisk | Extreme Pro 32Gb |
| 1 | USB cable | RND Components | Micro B to USB A |
| 1 | Station base | 3D printed | Black resin |
| 1 | Raspberry coupler | Laser cut | Acrylic 4mm |
| 2 | Arena's outer layers | Laser cut | Acrylic 1/16" |
| 1 | Arena's inner layer | Laser cut | Acrylic 1/16" |
| 1 | Prototyping board/PCB | Rademacher | 710-5 |
| 1 | Miniature Slide Switch | RND Components | 210-00585 |
| 1 | Trimmer Potentiometer | Bourns | 500 Ohms |
| 1 | Resistor | RND Components | 56 Ohms |
| 2 | Female-Female jumper wire | RND Components | BBFF-10-Q3RD |
| 4 | White LEDs | RND Components | 135-00164 |
| 4 | PCB pins angled | Prostar | RS-1X36-T1-7/3MM |
| 7 | Screws M2.5x12mm | Bossard | BN-610 |

### 4.7 2D pose estimation

DeepFly3D 2D poses were taken from [7]. OpenMonkeyStudio 2D poses were taken from [8]. CAPTURE 2D poses were taken from [39]. LocoMouse 2D poses were taken from [18]. See these publications for detailed 2D pose estimation procedures.

#### 4.7.1 2D pose estimation of freely behaving flies recorded in two views using a right-angle prism mirror

Data acquired from a single camera were split into ventral and side view images. We hand-annotated the location of all 30 leg joints (five joints per leg) on 640 images with a ventral view and up to 15 visible unilateral joints on 640 images of the side view. We used these manual annotations to train two separate DeepLabCut [6] 2D pose estimation networks (root-mean-squared errors for training and testing were 0.02 mm and 0.04 mm for ventral and side views, respectively). Whereas ventral view images could be used to predict 2D pose for all 30 leg joints, from the side view at most 15 joints were visible when the fly was parallel to the prism. Typically fewer keypoints were visible due to rotations of the fly within the enclosure. We removed images in which DeepLabCut incorrectly annotated keypoints as well as images in which flies were climbing the enclosure walls (thus exhibiting large yaw and roll orientation angles). To exclude these images, we ignored those with a confidence threshold below 0.95, and those for which the *x*-coordinate between the lateral and ventral views differed by more than 10 px.

#### 4.7.2 2D pose estimation of freely behaving flies recorded in one ventral view using a single camera

FlyLimbTracker data [20] was manually annotated because training a network to track only 100 frames would have been impractical. For newly acquired low-resolution ventral view single camera data, we trained a DeepLabCut [6] 2D pose estimation network. Due to the low resolution of images, the coxa-femur joints were not distinguishable, therefore, we treated the thorax-coxa and coxa-femur joints as a single entity. We manually annotated 160 images with the locations of four landmarks per leg: the thorax-coxa-femur entity, the femur-tibia joint, the tibia-tarsus joint, and the claw. We then trained a DeepLabCut network to predict the 2D coordinates of the 24 landmarks in the legs from the ventral view.

### 4.8 Training the LiftPose3D network

An important step in constructing LiftPose3D training data is to choose *r* root joints (see the specific use cases below for how these root joints were selected), and a target set corresponding to each root joint. The location of joints in the target set are predicted relative to the root joint to ensure translation invariance of the 2D poses.

The training dataset consisted of input-output pose pairs 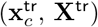 with dimensionality equal to the number of keypoints visible from a given camera *c* minus the number of root joints *r*, namely 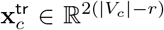 and 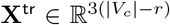. Then, the training data was standardized with respect to the mean and standard deviation of a given keypoint across all poses.

#### 4.8.1 Tethered *Drosophila melanogaster*

Of the 38 original keypoints in [7], here we focused on the 30 leg joints. Specifically, for each leg we estimated 3D position for the thorax-coxa, coxa-femur, femur-tibia, and tibia-tarsus joints and the tarsal tips (claws). Thus, the training data consisted of input-output coordinate pairs 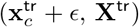 for 24 joints (30 minus six thorax-coxa root joints) from all cameras. Here 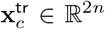 are 2D input joint keypoints acquired from camera *c* and **X**^tr^ ∈ R^3*n*^ are 3D ground truth coordinates obtained from DeepFly3D by triangulating 2D coordinates from all six cameras. Furthermore, *ϵ* ∈ ℝ^48^ is a small additive noise term, each with zero-mean Gaussian components. We found that the additive noise term stabilizes the network’s convergence during training (**Figure S2A**) and reduces uncertainty in lifted 3D joint positions. To maintain consistency for calculations of absolute error, triangulation was performed using the same set of 2D poses that were used to train the LiftPose3D network.

#### 4.8.2 Freely behaving macaque monkeys

The OpenMonkeyStudio dataset [8] consists of images of freely behaving monkeys inside a 2.45 × 2.45 × 2.75 m arena in which 62 cameras are equidistant horizontally at two heights along the arena perimeter. We extracted all five available experiments (7, 9, 9a, 9b and 11) for training and testing. Since 2D pose annotations were not available for all cameras, we augmented this dataset during training by projecting triangulated 3D poses onto cameras lacking 2D annotation using the provided camera matrix. For the available 2D annotations, we removed the fisheye lens-related distortions of 2D poses using the provided radial distortion parameters. We normalized each 2D pose to unit length, by dividing it by its Euclidean norm as well as the 3D pose with respect to bone lengths to reduce the large scale variability of the OpenMonkeyStudio annotations (animals ranged between 5.5 and 12 kg). Following the OpenMonkeyStudio convention, we set the neck joint as the root joint during training. We compare our absolute errors to the total body length, calculated as the sum of the mean lengths of the nose-neck, neck-hip, hip-knee, knee-foot joints pairs. Over multiple epochs, we observed rapid convergence of our trained network (**Figure S2B**).

#### 4.8.3 Freely behaving mice and *Drosophila* recorded from two views using a right-angle mirror

Freely behaving mouse data [18] consisted of recordings of animals traversing a 66.5 cm long, 4.5 cm wide, and 20 cm high glass corridor. A 45° mirror was used to obtain both ventral and side views with a single camera beneath the corridor. 2D keypoint positions were previously tracked using the LocoMouse software [18]. We considered six major keypoints—the four paws, the proximal tail, and the nose. Keypoint positions were taken relative to a virtual “root” keypoint placed on the ground midway between the nose and the tail.

For both the *Drosophila* and mouse datasets, side view keypoints distal to the camera were intermittently occluded by the animal’s body. Thus, taking a simplistic approach, after training with this unilateral ground truth data, lifting from the ventral view would only recover keypoints on the proximal half of the animal. We significantly modified data preprocessing to enable lifting across both the proximal and the occluded, distal side of the animal. Specifically, we registered all animals along the horizontal axis in the ventral view to generate ground truth data for all leg joints across time frames. Thus, although there is still only partial 3D pose ground truth for each image (for the proximal, fully visible half of the animal) we forced the lifting function *f* to predict the entire pose. This is possible because the realignment step masks from the network which data, among all of the input to *f*, are visible and contain 3D ground truth annotations.

Combining the proposed alignment and partial 3D pose supervision, the training dataset includes coordinate pairs 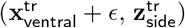, with *ϵ* as before, 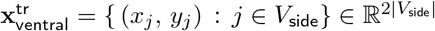 are the coordinates of DeepLabCut annotated 2D keypoints from the ventral viewpoint and 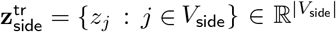 are the corresponding *z*-axis depth coordinates, for joints visible from the side view for a given frame. The networks for *Drosophila* and mouse training data converged within 30 and 10 training epochs (**Figure S2C,D**).

#### 4.8.4 Freely behaving rat in a naturalistic environment

The CAPTURE dataset contains recordings of freely behaving rats in a 2-foot diameter cylindrical arena tracked by six cameras. Motion capture markers on the animal are tracked using a commercial motion capture acquisition program [39] to obtain 2D poses. Out of 20 possible joints, we limited our scope to the 15 joints that were not redundant and provided most of the information about the animal pose. The dataset includes 4 experiments recording 3 rats from two different camera setups. Before using LiftPose3D, we removed the distortion from 2D poses using radial distortion parameters provided by the authors. The CAPTURE dataset has many missing 3D pose instances which we handle by not computing the loss corresponding to these keypoints during back-propagation. We selected the neck joint as the single root joint and predicted all of the other joints with respect to this root joint. We observed that LiftPose3D converged within 15 training epochs (**Figure S2E**).

#### 4.8.5 Freely behaving adult *Drosophila melanogaster* recorded from one ventral camera view

For both the newly acquired low-resolution and previously published high-resolution [20] images of freely behaving flies taken using one ventral view camera, we trained a LiftPose3D network on partial ground truth data acquired from the prism mirror system. For the high-resolution data, we considered the thorax-coxa joints as roots. For the low resolution data coxa-femur joints were imperceptible, allowing only 24 keypoints to be annotated. Hence, the thorax-coxa joints were selected as roots and we focused on predicting the relative location of the remaining mobile joints (18 keypoints) with respect to their associated root joints. The training dataset consisted of coordinate pairs 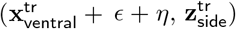 where 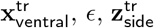 were chosen to represent the annotated ventral coordinates, joint-dependent noise and *z*-axis depth for the visible joints, as before. Meanwhile, *η* was a novel noise term, which we describe below.

The training and test data were augmented to accomplish domain adaptation: lifting new data with the prism system training data. First, for the low-resolution dataset, a zero-mean Gaussian noise term with a joint-independent standard deviation of 4 px, *η*, was added during training. The role of this noise term was to account for the keypoint position degeneracy inherent in the transformation from high-resolution prism training data to lower-resolution testing data. This term effectively coarse-grained the network’s spatial resolution, accounting for the 4-fold lower resolution of the low-resolution single camera ventral view system compared with the right-angle prism mirror system. For the high resolution dataset this noise term was set to zero.

Second, following training, we preprocessed the test data 2D poses derived from both the low- and high-resolution images by matching their data distributions to that of the prism-mirror dataset. To achieve this, we performed procrustes analysis to find the optimal affine transformation (rotation, translation and scaling) that maps the average root joint positions across poses in the test dataset to those in the prism-mirror dataset.

### 4.9 Deriving joint angles and performing error estimates

Consider three consecutive joints in the kinematic chain of one leg with coordinates **u, v, w**. Then, vectors **s**_1_ = **u**–**v** and **s**_2_ = **u**–**w** describe adjacent bones and their enclosed angle is found by the cosine rule, cos^*−*1^(**s**_1_ · **s**_2_*/*(||**s**_1_|| ||**s**_2_||)).

With the exception of the tarsus, the fly’s exoskeleton moves in a rigid manner. This permits the estimation of errors in the lifted joint angles based on fluctuations of predicted bone lengths. We assumed that **u, v, w** are drawn from independent Gaussian distributions centered around the estimated coordinate with standard deviation equal to the variation of the bone lengths ||**s**_1_|| and ||**s**_2_||. The distribution of joint angles for any given pose was estimated by Monte Carlo sampling (using 5 × 10^3^ samples) drawing one sample from each three distributions and then computing the corresponding joint angle by the cosine rule.

### 4.10 Code and data availability

The code can be installed as a pip package, see https://pypi.org/project/liftpose/, or downloaded at https://github.com/NeLy-EPFL/LiftPose3D.

The experimental data collected for this study can be downloaded at: https://drive.google.com/drive/folders/1qi8_c1YnlOzh7eWYXAG369iLtAS4iu1H?usp=sharing

## 5 Supplementary Notes

### 5.1 *Drosophila* LiftPose3D Station

#### Design and assembly

The LiftPose3D station’s main body was 3D printed using a Form2 printer and standard black resin (Formlabs, United States). A Raspberry Pi Zero W was fixed upside down onto the main body using a custom coupler and four screws. Two angled headers should be soldered to the Raspberry Pi ground (gnd) and 5V pins before placing the board onto the station. For the Raspberry Pi high-quality camera, the C-CS adapter was removed and the back focus adjustment ring was fully closed. Then, the 6 mm wide-angle lens was installed onto the camera. The camera and lens were screwed to a lasercut acrylic coupler with the cable connection facing the open side of the base, a flex cable was used to connect the camera to the Raspberry Pi board. We designed an illumination system consisting of four white LEDs, a switch to turn them on and off, and a potentiometer to control the light intensity. The circuit for controlling the LEDs was built on a prototyping board with prefabricated copper connections and we added extra connections with wires as shown in **Figure S3**. However, we also provide the files to manufacture a printed circuit board (PCB). The illumination module was screwed to the middle level of the base and two jumper wires were connected from the Raspberry Pi angled pins to the circuit pins considering the correct polarization, i.e., 5V to 5V and gnd to gnd. Finally, we used a square arena with three vertically stacked layers of 1/16” acrylic to hold behaving adult flies. The arena is 12 mm per side with rounded corners. These acrylic layers are fixed with the pillars on top of the base.

#### Raspberry Pi-Computer connection

We decided to establish a USB-Ethernet gadget mode connection to simplify the communication between our Raspberry Pi Zero and computer. This connection mode allowed us to power the Raspberry Pi, establish an SSH connection, and share the computer internet using one standard USB cable. However, any other connection mode can be used, including SSH through WiFi, or direct connection with a monitor, keyboard, and mouse to the Raspberry Pi board, as explained in the official Raspberry website.

We tested the USB-Ethernet gadget mode with a computer running Ubuntu 20.04, but different tutorials exist for running such a connection in iOS or Windows operating systems (OS). It is very important to use a standard USB cable and not an USB-OTG cable. First, a Raspberry Pi OS should be installed in a micro SD card using the Raspberry Pi Imager. We used the Raspberry Pi OS Lite version (Buster) with the Linux kernel 5.4.83.

After installing the OS, the SD card should be unplugged and plugged again into the PC. Now the card is mounted and we can acces the boot partition where some changes should be made.

First, the following lines should be appended to config.txt file to enable the OTG libraries on boot:

~~~
# Enable USB OTG like ethernet
dtoverlay=dwc2
~~~

Then, an empty file called ssh (without any extension) should be created using, e.g., vim, vi, or touch. Finally, modify the cmdline.txt file by adding the following line after the word “rootwait” (add a space at the beginning and the end of the added text):

~~~
modules-load=dwc2,g_ether
~~~

Now that the initial configuration is completed, the SD card should be ejected from the computer and inserted into the Raspberry board. Then, connect the USB cable to the USB port labeled “USB”, not the one labeled “PWR”. Booting the first time lasts around 60-90 s, afterwards it will be faster. The Raspberry Pi will be recognized by Ubuntu as a new Ethernet network connection. However, to enable the connection, it has to be edited to set the connection method to “Link-Local Only” in the IPv4 tab. The ssh tunneling is established and the Raspberry can be accessed using:

~~~
ssh pi@raspberrypi.local
~~~

By default the ssh password is “raspberry”, however, it can be easily changed. To share the internet connection from the Ubuntu computer, the ethernet connection should be disconnected in the networks manager (do not disconnect the USB cable) and the connection method should be changed to “Shared to other computers”. Establish again the ethernet connection and now the Raspberry Pi will have internet after you ssh onto it.

For now the Raspberry Pi will choose a random ID and MAC-address after every restart/reboot. To fix that, edit again the cmdline.txt file on the boot partition by adding the following line at the end:

~~~
g_ether.host_addr=xx:xx:xx:xx:xx:xx
~~~

The host address should be taken from the Ubuntu computer and can be obtained by running the command `ifconfig` in a terminal. The last thing to do is to assign a static IP address to the Raspberry. To do that, add the following lines to the file “/etc/dhcpcd.conf”:

~~~
interface usb0
static ip_address=10.xx.xx.xx
static routers=10.xx.xx.xx
~~~

The static IP address and routers are then obtained from the Raspberry and the Ubuntu computer, respectively, by running the command `ifconfig`. The whole configuration above should be done just once. At this point the connection between the Raspberry and the computer will be established automatically every time the USB cable is used.

#### Image acquisition

To set up the acquisition software in the LiftPose3D station, first python 3 and pip3 should be installed on the Raspberry Pi Zero:

~~~
$ sudo apt-get update
$ sudo apt-get upgrade
$ sudo apt-get install python3-dev
$ wget https://bootstrap.pypa.io/get-pip.py
$ sudo python3 get-pip.py
~~~

Then the Raspberry Pi camera should be enabled by running `$ sudo raspi-config`, and selecting the corresponding option. The Raspberry should be rebooted after enabling the camera module. Finally, the package piCamera should be installed by running the command:

~~~
$ pip install “picamera[array]”.
~~~

The script *capture fast.py* should be copied in the Raspberry Pi and it can be run with the command:

~~~
$ python3 capture_fast.py imgsFolder
~~~

The script is a customized version of an example (Advance recipe 4.7) found in the piCamera package documentation. It will capture images for 30 s by default at 80 fps with an exposure time fixed at 2 ms. These images will be stored in a directory named `imgsFolder`. The recording duration, framerate, and exposure time can be modified directly in the program, however, it is not recommended to change either the framerate or the exposure time since it would change the illumination and sharpness of the images.

#### Image preprocessing

When the images are captured they are stored onto the Raspberry SD card, however, we strongly recommend moving them to another computer with larger capacity as soon as they are taken. A preprocessing stage should be completed before lifting the fly’s pose. This procedure consists of cropping the fly from every frame and registering these crops along the experiment aligning the fly facing up. This processing is performed by the program *crop flies.py*, using the following pseudocode:

1. Read frames.
2. Segment fly’s body based on color.
3. Binarize image.
4. Fit ellipse around the fly’s body.
5. Get crop of 290×290 pixels around the ellipse centroid.
6. Rotate crop based on the ellipse orientation to register images.
7. Check for head or wings at the top of the crop.
8. Rotate crops if wings detected on top in more than 50% of the frames.
9. Write video with cropped fly.

## 7 Supplementary Figures

**Figure S1:**
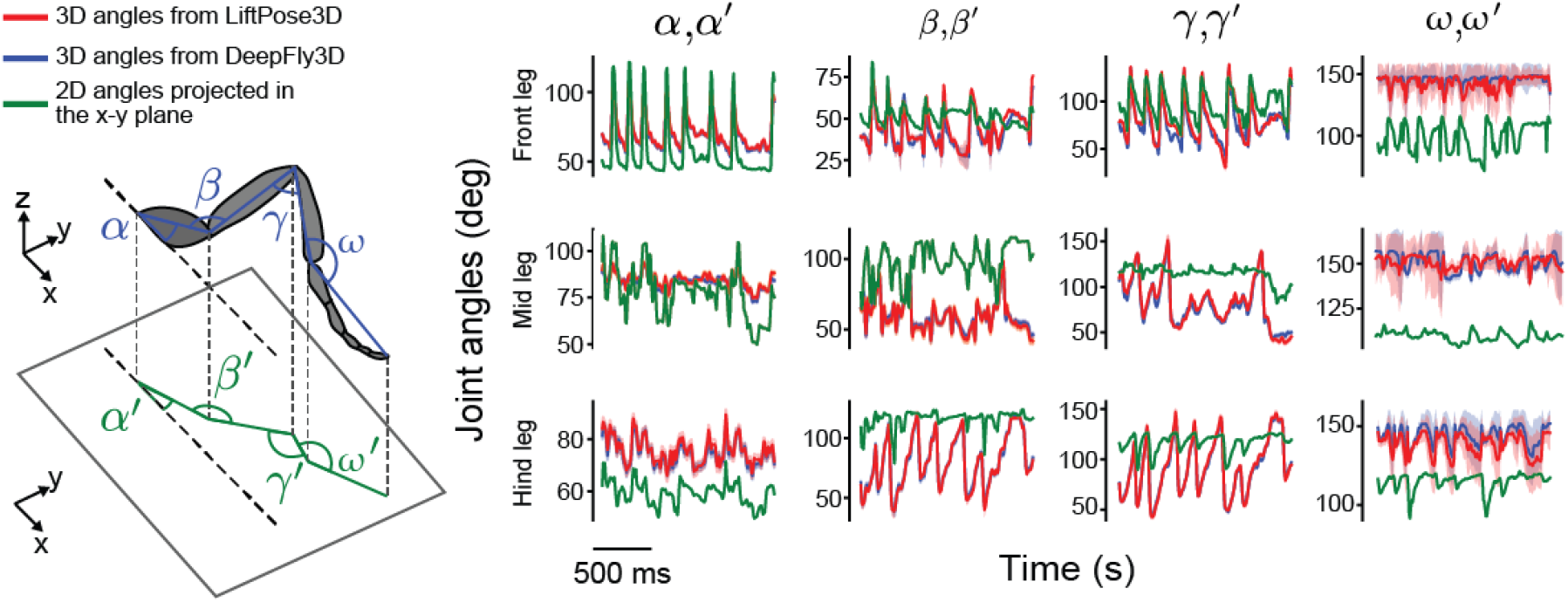
Joint angles resulting from lifting compared with 3D triangulated ground truth and 2D projections. Joint angles *α, β, γ*, and *ω* for the front, mid, and hind left legs during forward walking. Shown are angles computed from 3D triangulation using DeepFly3D (blue), LiftPose3D predictions (red), and ventral 2D projections *α*′, *β*′, *γ*, and *ω* (green). The mean (solid lines) and standard deviation of joint error distributions (transparency) are shown. Joint angles were computed by Monte Carlo sampling and errors were computed by taking the fluctuation in bone lengths.

**Figure S2:**
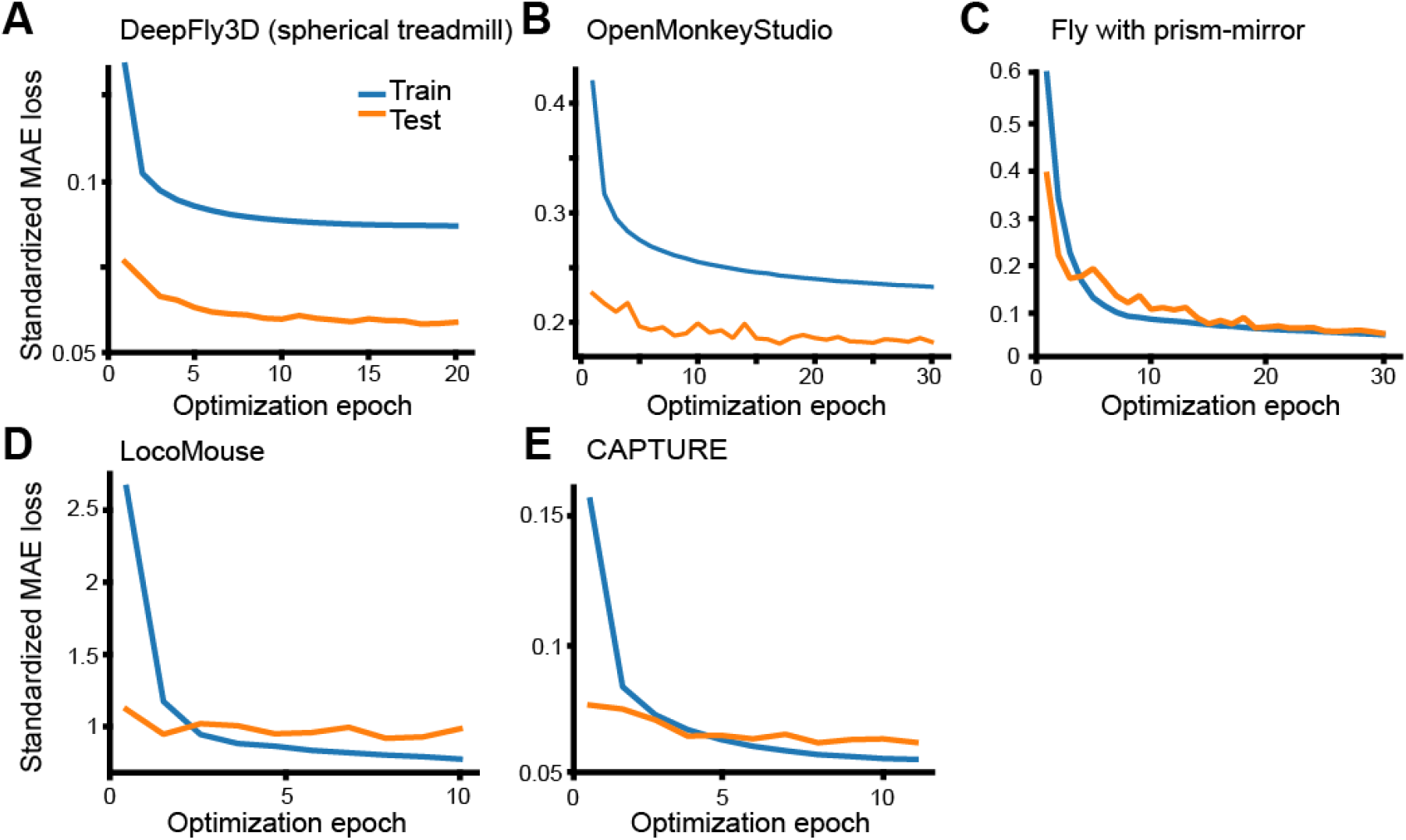
Training and test loss convergence of the LiftPose3D network applied to a variety of datasets. Shown are the absolute test errors of LiftPose3D for all joints as a function of optimization epoch. Note that the test error is sometimes lower than the training error because we do not apply dropout at test time. **A** Two-camera data of *Drosophila* on a spherical treadmill (each color denotes a different pair of diametrically opposed cameras). **B** OpenMonkeyStudio dataset (each color denotes a different training run). **C** Single-camera data of *Drosophila* behaving freely in the right-angle prism mirror system, **D** LocoMouse dataset. **E** CAPTURE dataset.

**Figure S3:**
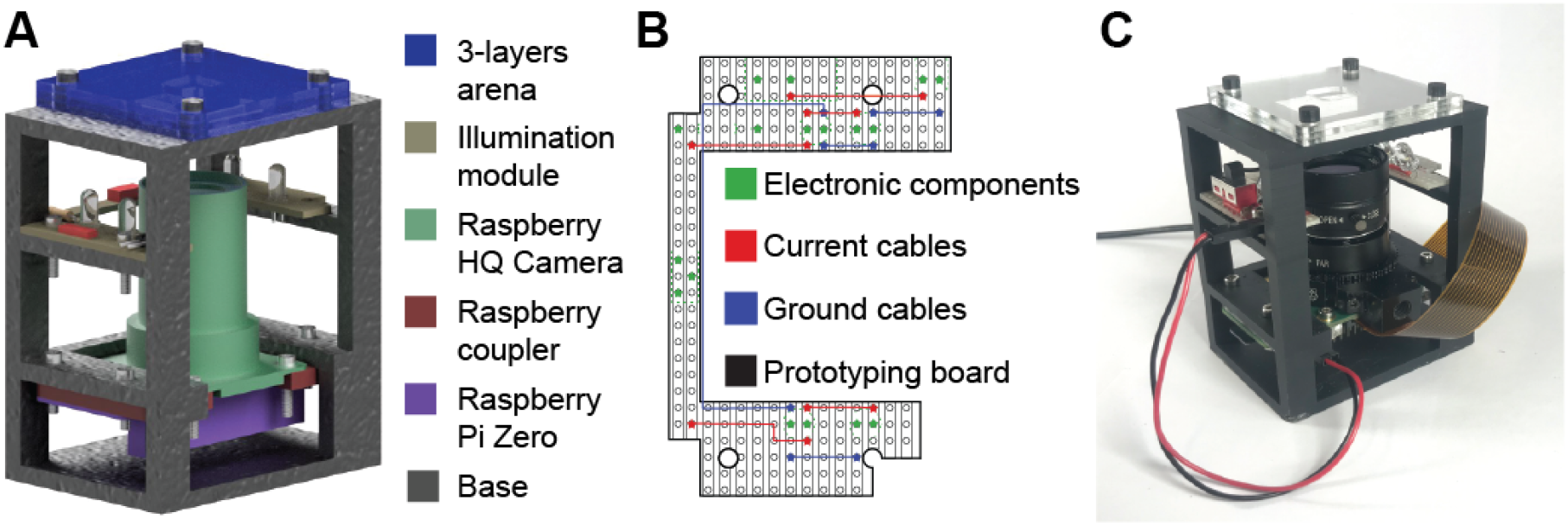
*Drosophila* LiftPose3D station. **A** CAD drawing of the LiftPose3D station indicating major components (color-coded). **B** Electronic connections included in a prefabricated prototyping board for the illumination module. **C** Photo of the LiftPose3D station.

## 8 Supplementary Videos

**Video 1: 3D pose lifting for backwards walking in tethered *Drosophila* obtained from two side cameras**. Videos obtained from cameras 2 **(top-left)** and 5 **(bottom-left)**. DeepFly3D-derived 2D poses are superimposed. Orange circle indicates that the optogenetic stimulation LED light is on, activating MDNs to elicit backward walking. **(right)** 3D poses obtained by triangulating six camera views using DeepFly3D (solid lines), or lifting two camera views using LiftPose3D (dashed lines).

https://www.dropbox.com/s/e1dpxqf23epxtg2/video_1.mp4?dl=0

**Video 2: 3D pose lifting for antennal grooming in tethered *Drosophila* obtained from two side cameras**. Videos obtained from cameras 2 **(top-left)** and 5 **(bottom-left)**. DeepFly3D-derived 2D poses are superimposed. Orange circle indicates that the optogenetic stimulation LED light is on, activating aDNs to elicit antennal grooming. **(right)** 3D poses obtained by triangulating six camera views using DeepFly3D (solid lines), or lifting two camera views using LiftPose3D (dashed lines).

https://www.dropbox.com/s/fzvru50z43a9t9t/video_2.mp4?dl=0

**Video 3: 3D pose lifting for irregular spontaneous limb movements in tethered *Drosophila* obtained from two side cameras**. Videos obtained from cameras 2 **(top-left)** and 5 **(bottom-left)**. DeepFly3D-derived 2D poses are superimposed. **(right)** 3D poses obtained by triangulating six camera views using DeepFly3D (solid lines), or lifting two camera views using LiftPose3D (dashed lines).

https://www.dropbox.com/s/5qbdiq9fdtlkgdo/video_3.mp4?dl=0

**Video 4: 3D pose lifting of previously published OpenMonkeyStudio dataset of a freely moving macaque [8] (left)** Single image drawn randomly from one of 62 cameras. **(middle)** Ground truth 3D poses based on triangulation of 2D poses from up to 62 cameras (solid lines), or lifting from a single camera view using LiftPose3D (dashed lines). **(right)** Error distribution across the 62 cameras for a given pose. Camera locations (circles) are color-coded by error. Gray circles denote cameras for which an image was not available. Green circle denotes the camera from which the image was used.

https://www.dropbox.com/s/mfe32jnen9oo6w8/video_4.mp4?dl=0

**Video 5: 3D pose lifting of freely behaving *Drosophila* when triangulation is only partially possible**. Single camera images of the ventral **(top-left)** and side **(bottom-left)** views. DeepLabCut-derived 2D poses are superimposed. **(right)** 3D poses obtained by triangulating partially available multi-view 2D poses (solid lines), or by lifting the ventral 2D pose using LiftPose3D (dashed lines).

https://www.dropbox.com/s/1cd36l55kda89pq/video_5.mp4?dl=0

**Video 6: 3D pose lifting of freely behaving mice when triangulation is only partially possible**. Side **(top-left)** and ventral **(bottom-left)** views of a freely walking mouse. Superimposed are keypoints on the paws, mouth, and proximal tail tracked using the LocoMouse software (blue circles). Using only the ventral view 2D pose, a trained LiftPose3D network can accurately track keypoints in the side view (orange circles).

https://www.dropbox.com/s/jh2xaqfmf2wmd8p/video_6.mp4?dl=0

**Video 7: 3D pose lifting for freely behaving rats in a naturalistic arena (left)** Ground truth 3D poses triangulated from six cameras (solid lines) superimposed with LiftPose3D’s predictions using 2D poses from one camera (dashed lines). **(right)** Images from one camera with 2D poses acquired using CAPTURE are superimposed.

https://www.dropbox.com/s/1awphk5gfc2u9pc/video_7.mp4?dl=0

**Video 8: 3D pose lifting for low-resolution videos of freely behaving flies when triangulation is impossible. (top)** Three freely behaving *Drosophila* in a rounded square arena and recorded ventrally using a single low-resolution camera. Of these, fly 0 is tracked, cropped, and rotated leftward. Superimposed are 2D poses for 24 visible joints. (bottom) 3D poses lifted from ventral view 2D poses (*x*− *y* plane) permit analysis of leg kinematics in the otherwise unobserved *x* −*z* plane.

https://www.dropbox.com/s/7we9lcp2n74c838/video_8.mp4?dl=0

**Video 9: 3D pose lifting of previously published ventral view videos of freely behaving flies when triangulation is impossible. (top)** Video from [20] of a freely behaving fly within a pill-shaped arena and recorded ventrally using a single high-resolution camera. **(bottom-left)** Following tracking, a region-of-interest containing the fly was cropped and rotated to maintain a leftward orientation. Superimposed are 2D poses estimated for 24 visible joints. **(bottom-middle)** 3D poses obtained by lifting ventral view 2D poses. **(bottom-right)** 3D poses lifted from ventral view 2D poses (top) permit analysis of leg kinematics in the otherwise unobserved *x* −*z* plane (bottom). https://www.dropbox.com/s/2tylyqcnqgdq4qc/video_9.mp4?dl=0

**Video 10: 3D pose lifting of data from the *Drosophila* LiftPose3D station (left)** Video of a freely behaving fly in the LiftPose3D station arena. **(middle)** Cropped video around the centroid of the tracked fly, superimposed with 2D pose predictions. **(right)** Lifted 3D poses obtained using ventral 2D poses.

https://www.dropbox.com/s/esnwx0we5itteb6/video_10.mp4?dl=0

## 10 Acknowledgments

We thank the Ölvecky lab for providing us with the CAPTURE dataset. We thank Megan Carey (Champalimaud Centre for the Unknown, Lisbon, Portugal) for the LocoMouse dataset.

## 11 Author Contributions

A.G. - Conceptualization, methodology, software, hardware (*Drosophila* prism mirror system), formal analysis, data curation, writing—original draft, writing—review & editing.

S.G. - Conceptualization, methodology, software, formal analysis, data curation, writing—original draft, writing—review & editing.

V.L.R. - Software and hardware (LiftPose3D station, low-resolution *Drosophila* ventral view system), data curation, writing—review & editing.

M.A. - Methodology, software (LiftPose3D), preliminary analysis of DeepFly3D dataset, data curation, writing—review & editing.

D.M. - Investigation (low-resolution *Drosophila* experiments), writing—review & editing.

H.R. - Conceptualization, writing—review & editing.

P.F. - Writing—review & editing, funding acquisition.

P.R. - Conceptualization, hardware (*Drosophila* prism mirror system), resources, writing—original draft, writing—review & editing, supervision, project administration, funding acquisition.

## References

[1] Dombeck, D. A., Khabbaz, A. N., Collman, F., Adelman, T. L. & Tank, D. W. Imaging large-scale neural activity with cellular resolution in awake, mobile mice. Neuron 56, 43–57 (2007).

[2] Seelig, J. D. et al. Two-photon calcium imaging from head-fixed Drosophila during optomotor walking behavior. Nature Methods 7, 535–540 (2010).

[3] Churchland, M. M. et al. Neural population dynamics during reaching. Nature 487, 51–56 (2012).

[4] Chen C. L., Hermans L. et al. Imaging neural activity in the ventral nerve cord of behaving adult drosophila. Nature communications 9, 4390 (2018).

[5] Pereira, T. D. et al. Fast animal pose estimation using deep neural networks. Nature Methods 16, 117–125 (2019).

[6] Mathis, A. et al. DeepLabCut: markerless pose estimation of user-defined body parts with deep learning. Nature neuroscience 21, 1281–1289 (2018).

[7] Günel, S. et al. DeepFly3D, a deep learning-based approach for 3D limb and appendage tracking in tethered, adult Drosophila. eLife 8, 3686 (2019).

[8] Bala, P. C. et al. OpenMonkeyStudio: Automated markerless pose estimation in freely moving macaques. bioRxiv (2020).

[9] Newell, A., Yang, K. & Deng, J. Stacked hourglass networks for human pose estimation. In European Conference on Computer Vision (ECCV) (2016).

[10] Graving, J. M. et al. Deepposekit, a software toolkit for fast and robust animal pose estimation using deep learning. eLife 8, e47994 (2019).

[11] Fang, H.-S., Xie, S., Tai, Y.-W. & Lu, C. RMPE: Regional multi-person pose estimation. In IEEE International Conferene on Computer Vision (ICCV) (2017).

[12] Wei, S.-E., Ramakrishna, V., Kanade, T. & Sheikh, Y. Convolutional pose machines. In IEEE Conference on Computer Vision and Pattern Recognition (CVPR) (2016).

[13] Cao, Z., Simon, T., Wei, S.-E. & Sheikh, Y. Realtime multi-person 2D pose estimation using part affinity fields. In IEEE Conference on Computer Vision and Pattern Recognition (CVPR) (2017).

[14] Hartley, R. & Zisserman, A. Multiple View Geometry in Computer Vision (Cambridge University Press, USA, 2003), 2 edn.

[15] Karashchuk, P. et al. Anipose: a toolkit for robust markerless 3D pose estimation. bioRxiv (2020).

[16] Nath, T., Mathis, A., Chen, A. C., Bethge, M. & Mathis, M. W. Using DeepLabCut for 3D markerless pose estimation across species and behaviors. Nature Protocols 14, 2152–2176 (2019).

[17] Gaudry, Q., Hong, E. J., Kain, J., de Bivort, B. L. & Wilson, R. I. Asymmetric neurotransmitter release enables rapid odour lateralization in Drosophila. Nature 493, 424–428 (2013).

[18] Machado, A. S., Darmohray, D. M., Fayad, J., Marques, H. G. & Carey, M. R. A quantitative framework for whole-body coordination reveals specific deficits in freely walking ataxic mice. eLife 4, e07892 (2015).

[19] Isakov, A. et al. Recovery of locomotion after injury in Drosophila melanogaster depends on proprioception. Journal of Experimental Biology 219, 1760–1771 (2016).

[20] Uhlmann, V., Ramdya, P., Delgado-Gonzalo, R., Benton, R. & Unser, M. Flylimbtracker: An active contour based approach for leg segment tracking in unmarked, freely behaving Drosophila. PLoS One 12, e0173433 (2017).

[21] DeAngelis, B. D., Zavatone-Veth, J. A. & Clark, D. A. The manifold structure of limb coordination in walking Drosophila. eLife 8, 137 (2019).

[22] Lee, H.-J. & Chen, Z. Determination of 3D human body postures from a single view. Computer Vision, Graphics, and Image Processing 30, 148–168 (1985).

[23] Taylor, C. J. Reconstruction of articulated objects from point correspondences in a single un-calibrated image. In IEEE Conference on Computer Vision and Pattern Recognition (CVPR) (2000).

[24] Chen, C. & Ramanan, D. 3D human pose estimation = 2D pose estimation + matching. In IEEE Conference on Computer Vision and Pattern Recognition (CVPR) (2017).

[25] Gupta, A., Martinez, J., Little, J. J. & Woodham, R. J. 3D pose from motion for cross-view action recognition via non-linear circulant temporal encoding. In IEEE Conference on Computer Vision and Pattern Recognition (CVPR) (2014).

[26] Sun, J. J. et al. View-invariant probabilistic embedding for human pose. Preprint at https://arxiv.org/abs/1912.01001 (2019).

[27] Nibali, A., He, Z., Morgan, S. & Prendergast, L. 3D human pose estimation with 2D marginal heatmaps. In IEEE Winter Conference on Applications of Computer Vision (WACV) (2019).

[28] Zhao, L., Peng, X., Tian, Y., Kapadia, M. & Metaxas, D. N. Semantic graph convolutional networks for 3D human pose regression. In IEEE Conference on Computer Vision and Pattern Recognition (CVPR) (2019).

[29] Iskakov, K., Burkov, E., Lempitsky, V. & Malkov, Y. Learnable triangulation of human pose. In International Conference on Computer Vision (ICCV) (2019).

[30] Kanazawa, A., Zhang, J. Y., Felsen, P. & Malik, J. Learning 3D human dynamics from video. In IEEE Conference on Computer Vision and Pattern Recognition (CVPR) (2019).

[31] Mehta, D. et al. XNect: Real-time multi-person 3D motion capture with a single RGB camera. In ACM Transactions on Graphics (2020).

[32] Rematas, K., Nguyen, C., Ritschel, T., Fritz, M. & Tuytelaars, T. Novel views of objects from a single image. Preprint at https://arxiv.org/pdf/1602.00328 (2016).

[33] Rhodin, H., Constantin, V., Katircioglu, I., Salzmann, M. & Fua, P. Neural scene decomposition for multi-person motion capture. In IEEE Conference on Computer Vision and Pattern Recognition (CVPR) (2019).

[34] Martinez, J., Hossain, R., Romero, J. & Little, J. J. A simple yet effective baseline for 3D human pose estimation. In IEEE International Conference on Computer Vision (ICCV) (2017).

[35] Pavllo, D., Feichtenhofer, C., Grangier, D. & Auli, M. 3D human pose estimation in video with temporal convolutions and semi-supervised training. In IEEE Conference on Computer Vision and Pattern Recognition (CVPR) (2019).

[36] Liu, J., Guang, Y. & Rojas, J. GAST-Net: Graph attention spatio-temporal convolutional networks for 3D human pose estimation in video. Preprint at https://arxiv.org/abs/2003.14179 (2020).

[37] Cai, Y. et al. Exploiting spatial-temporal relationships for 3D pose estimation via graph convolutional networks. In IEEE International Conference on Computer Vision (ICCV) (2019).

[38] Yiannakides, A., Aristidou, A. & Chrysanthou, Y. Real-time 3D human pose and motion reconstruction from monocular rgb videos. Comput. Animat. Virtual Worlds 30, 1–12 (2019).

[39] Marshall, J. D. et al. Continuous whole-body 3D kinematic recordings across the rodent behavioral repertoire. Neuron 109, 420–437.e8 (2021).

[40] Srivastava, N., Hinton, G., Krizhevsky, A., Sutskever, I. & Salakhutdinov, R. Dropout: a simple way to prevent neural networks from overfitting. The Journal of Machine Learning Research 15, 1929–1958 (2014).

[41] Wandt, B., Rudolph, M., Zell, P., Rhodin, H. & Rosenhahn, B. CanonPose: Self-supervised monocular 3D human pose estimation in the wild. Preprint at https://arxiv.org/abs/2011.14679 (2020).

[42] Wei, S., Ramakrishna, V., Kanade, T. & Sheikh, Y. Convolutional pose machines. In IEEE Conference on Computer Vision and Pattern Recognition (CVPR) (2016).

[43] Card, G. & Dickinson, M. H. Visually mediated motor planning in the escape response of Drosophila. Current Biology 18, 1300–1307 (2008).

[44] Wosnitza, A., Bockemühl, T., Dübbert, M., Scholz, H. & Büschges, A. Inter-leg coordination in the control of walking speed in Drosophila. Journal of experimental biology 216, 480–491 (2013).

[45] Cao, J. et al. Cross-domain adaptation for animal pose estimation. Preprint at https://arxiv.org/abs/1908.05806 (2019).

[46] Sanakoyeu, A., Khalidov, V., McCarthy, M. S., Vedaldi, A. & Neverova, N. Transferring Dense Pose to Proximal Animal Classes. In IEEE Conference on Computer Vision and Pattern Recognition (CVPR) (2020).

[47] De Bono, M. & Bargmann, C. I. Natural variation in a neuropeptide y receptor homolog modifies social behavior and food response in c. elegans. Cell 94, 679–689 (1998).

[48] Budick, S. A. & O’Malley, D. M. Locomotor repertoire of the larval zebrafish: swimming, turning and prey capture. Journal of Experimental Biology 203, 2565–2579 (2000).

[49] Louis, M., Huber, T., Benton, R., Sakmar, T. P. & Vosshall, L. B. Bilateral olfactory sensory input enhances chemotaxis behavior. Nature neuroscience 11, 187–199 (2008).

[50] Strauss, R. & Heisenberg, M. Coordination of legs during straight walking and turning in Drosophila melanogaster. Journal of Comparative Physiology A 167, 403–412 (1990).

[51] Clarke, K. & Still, J. Gait analysis in the mouse. Physiology & behavior 66, 723–729 (1999).

[52] Wiltschko, A. B. et al. Mapping sub-second structure in mouse behavior. Neuron 88, 1121–1135 (2015).

[53] Hong, W. et al. Automated measurement of mouse social behaviors using depth sensing, video tracking, and machine learning. Proceedings of the National Academy of Sciences 112, E5351–E5360 (2015).

[54] Mendes, C. S., Bartos, I., Akay, T., Márka, S. & Mann, R. S. Quantification of gait parameters in freely walking wild type and sensory deprived Drosophila melanogaster. elife 2, 231 (2013).

[55] Feng, K. et al. Distributed control of motor circuits for backward walking in drosophila. Nature communications 11, 1–17 (2020).

[56] Alp Güler, R., Neverova, N. & Kokkinos, I. Densepose: Dense human pose estimation in the wild. In IEEE Conference on Computer Vision and Pattern Recognition (CVPR) (2018).

[57] Güler, R. A. & Kokkinos, I. Holopose: Holistic 3D human reconstruction in-the-wild. In IEEE Conference on Computer Vision and Pattern Recognition (CVPR) (2019).

[58] Loper, M., Mahmood, N., Romero, J., Pons-Moll, G. & Black, M. J. SMPL: A skinned multiperson linear model. ACM Trans. Graphics (Proc. SIGGRAPH Asia) 34, 248:1–248:16 (2015).

[59] Zhang, J. Y., Felsen, P., Kanazawa, A. & Malik, J. Predicting 3D human dynamics from video. In IEEE International Conference on Computer Vision (ICCV) (2019).

[60] Zuffi, S., Kanazawa, A., Berger-Wolf, T. & Black, M. J. Three-d safari: Learning to estimate zebra pose, shape, and texture from images ”in the wild”. In IEEE International Conferene on Computer Vision (ICCV) (2019).

[61] Nair, V. & Hinton, G. E. Rectified linear units improve restricted boltzmann machines. In International Conference on Machine Learning (ICML), 807–814 (2010).

[62] He, K., Zhang, X., Ren, S. & Sun, J. Deep residual learning for image recognition. In IEEE Conference on Computer Vision and Pattern Recognition (CVPR) (2016).

[63] Kingma, D. P. & Ba, J. Adam: A method for stochastic optimization. Preprint at https://arxiv.org/abs/1412.6980 (2014).

[64] Ioffe, S. & Szegedy, C. Batch normalization: Accelerating deep network training by reducing internal covariate shift. In International conference on machine learning, 448–456 (PMLR, 2015).

[65] Sridhar, V. H., Roche, D. G. & Gingins, S. Tracktor: Image-based automated tracking of animal movement and behaviour. Methods in Ecology and Evolution 10, 815–820 (2019).

